# Linear-time cluster ensembles of large-scale single-cell RNA-seq and multimodal data

**DOI:** 10.1101/2020.06.15.151910

**Authors:** Van Hoan Do, Francisca Rojas Ringeling, Stefan Canzar

## Abstract

A fundamental task in single-cell RNA-seq (scRNA-seq) analysis is the identification of transcriptionally distinct groups of cells. Numerous methods have been proposed for this problem, with a recent focus on methods for the cluster analysis of ultra-large scRNA-seq data sets produced by droplet-based sequencing technologies. Most existing methods rely on a sampling step to bridge the gap between algorithm scalability and volume of the data. Ignoring large parts of the data, however, often yields inaccurate groupings of cells and risks overlooking rare cell types. We propose method *Specter* that adopts and extends recent algorithmic advances in (fast) spectral clustering. In contrast to methods that cluster a (random) subsample of the data, we adopt the idea of landmarks that are used to create a sparse representation of the *full* data from which a spectral embedding can then be computed in linear time. We exploit Specter’s speed in a cluster ensemble scheme that achieves a substantial improvement in accuracy over existing methods and that is sensitive to rare cell types. Its linear time complexity allows Specter to scale to millions of cells and leads to fast computation times in practice. Furthermore, on CITE-seq data that simultaneously measures gene and protein marker expression we demonstrate that Specter is able to utilize multimodal omics measurements to resolve subtle transcriptomic differences between subpopulations of cells. Specter is open source and available at https://github.com/canzarlab/Specter.

## Introduction

Single-cell RNA sequencing (scRNA-seq) has dramatically increased the resolution at which important questions in cell biology can be addressed. It has helped to identify novel cell types based on commonalities and differences in genome-wide expression patterns, reconstruct the heterogeneous composition of cell populations in tumors and their microenvironment, and unveil regulatory programs that govern the dynamic changes in gene expression along developmental trajectories.

One of the most fundamental computational tasks in the context of scRNA-seq analysis is the identification of groups of cells that are similar in their expression patterns, i.e. their transcriptomes, and which are at the same time distinct from other cells. Conceptually similar problems have been studied in anthropology (Driver and Kroeber 1932) and psychology (Zubin 1938) almost a century ago and since then this so-called cluster analysis has become one of the most well-studied problems in unsupervised machine learning. Numerous methods have been proposed for clustering scRNA-seq data sets (Duò et al. 2018; Tian et al. 2019), with Seurat (Satija et al. 2015) and its underlying Louvain clustering algorithm (Blondel et al. 2008) being arguably the most widely used one. More recently, attempts have been made to design algorithms for the analysis of ultra-large scRNA-seq data sets, owing to the ever-increasing throughput of droplet-based sequencing technologies that allow to profile genome-wide expression for hundreds of thousands of cells at once. At the heart of such methods often lies a sampling technique that reduces the size of the data analyzed by a clustering algorithm. Cluster labels of cells in this so-called sketch are subsequently transferred to the remaining cells using, e.g., a nearest neighbor algorithm. dropClust (Sinha et al. 2018), for example, includes a structure preserving sampling step, but initially picks a small set of cells simply at random. Similarly, Seurat applies random subsampling prior to its nearest neighbor search.

The quality of the final clustering, however, strongly depends on how well the data sketch represents the overall cluster structure and how accurate the cluster labels of cells in the sketch can be inferred from incomplete data. Inaccurate labels of subsampled cells will likely lead to an inaccurate labeling of the full data. In addition, sampling cells proportional to their abundance might render rare cell types invisible to the algorithm. Geometric sketching was therefore recently proposed as an alternative sampling method that selects cells according to the transcriptomic space they occupy rather than their abundance. Nevertheless, labels need to be inferred from partial data.

Spectral methods for clustering have been applied with great success in many areas such as computer vision, robotics, and bioinformatics. They make few assumptions on cluster shapes and are able to detect clusters that form non-convex regions. On a variety of data types, this flexibility has allowed spectral clustering methods to produce more accurate clusterings than competing methods (Shi and Malik 2000). The high computational complexity, however, renders its application to large-scale problems infeasible. For *n* data points, spectral clustering computes eigenvectors of a *n* × *n* affinity matrix, which incurs a computational cost of 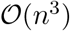. For scRNA-seq data sets with *n* in the order of ten thousands up to the millions this presents a prohibitive cost which has thus prevented the application of spectral clustering to large-scale single-cell data sets.

Furthermore, spectral clustering methods are sensitive to the right choice of parameters used to model the similarity between data points (Luxburg 2007), i.e. RNA expression measurements of single cells. Data sets derived from different biological samples exhibiting different cell population structures obtained using different sequencing technologies typically require a different set of parameter values to achieve accurate clustering results.

We introduce a new method, Specter, which addresses the challenges of computational complexity and parameter sensitivity to allow a tailored version of spectral clustering to be utilized in the analysis of large scRNA-seq data sets. In contrast to methods that cluster only a (random) subsample of the data, Specter takes a fundamentally different approach that avoids learning from unlabeled partial data. We adopt the idea of landmarks (Cai and Chen 2011), a random sample of cells that are used to create a sparse representation of the *full* data from which a spectral embedding can then be computed in linear time. We utilize the speed of this approach to systematically explore the parameter space by a co-association based consensus clustering scheme, also known in literature as cluster ensembles (Strehl and Ghosh 2003): Instead of picking one set of parameters, in Specter we explore different choices of parameters and reconcile the resulting clustering information into a single (consensus) clustering. In addition to aggregating clusterings obtained in different runs of the algorithm on the same data, consensus clustering can also be used to reconcile clusterings of cells based on different molecular features. Recent technological advances have allowed to simultaneously measure multiple modalities of single cells (Zhu et al. 2020). CITE-seq (Stoeckius et al. 2017), for example, measure both gene expression and surface protein levels of individual cells and Specter’s consensus clustering scheme can help to resolve subpopulations of cells that cannot accurately be distinguished based on transcriptomic differences alone. We combine consensus clustering with a novel *selective sampling* strategy that utilizes clustering information obtained from the full data set to achieve overall linear time complexity. Finally, we transfer cluster labels to the remaining cells using supervised *k*-nearest neighbors classification. We provide an overview of the approach in Figure 1.

**Figure 1:**
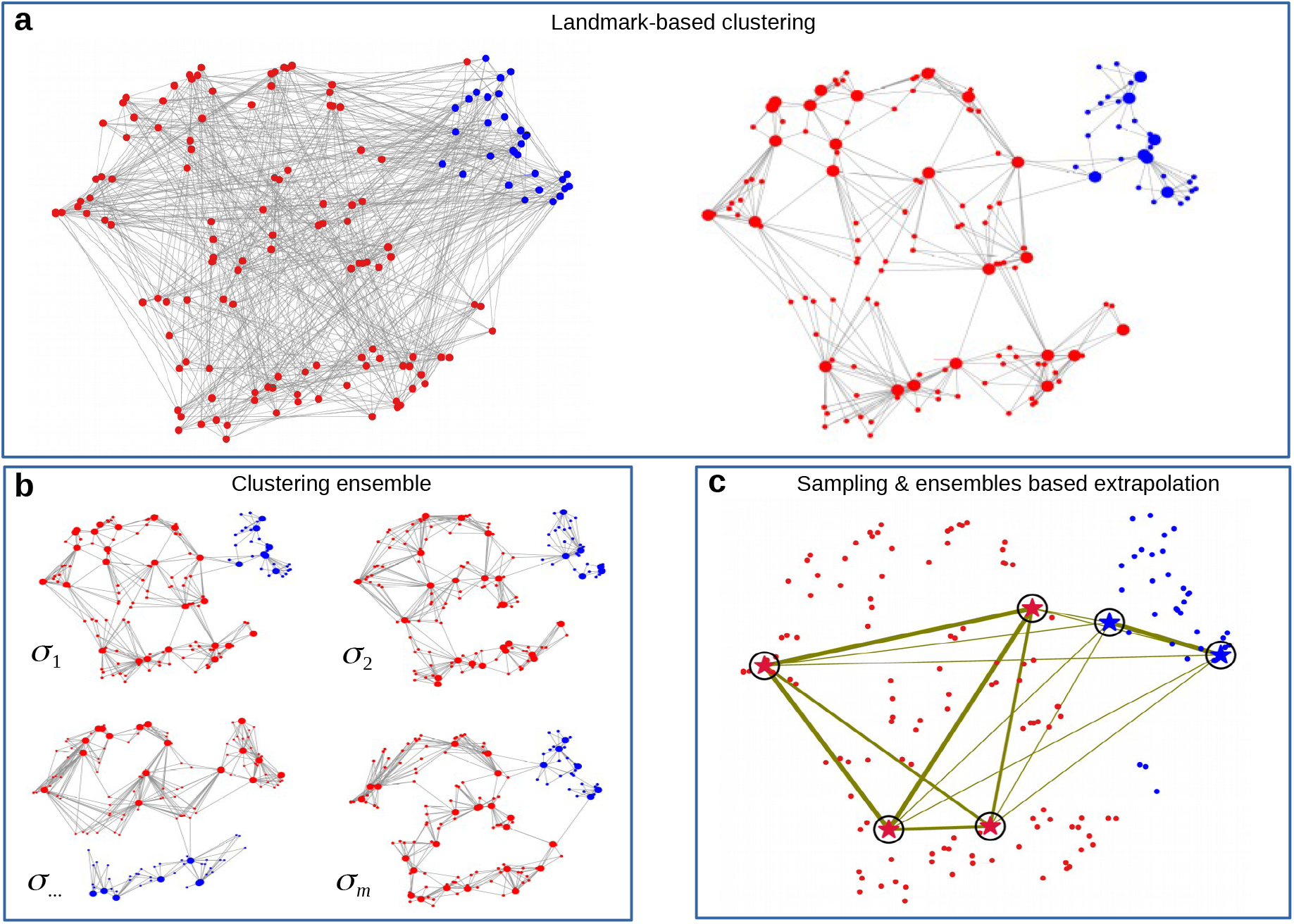
Overview of Specter. Illustrations are based on t-SNE visualizations of a random subsample of scRNA-seq data by Grün et al. 2016. (**a**) Standard spectral clustering constructs an affinity matrix that captures (transcriptional) similarities between all pairs of cells (left) which renders its eigen decomposition prohibitively expensive for large data sets. In contrast (right), describing each cell (small circles) with respect to its nearby *landmarks* (big circles) that were initially selected as the means computed by *k*-means clustering, creates a sparse representation of the full data that dramatically speeds up the computation of a spectral embedding. Cell are colored to distinguish sorted hematopoetic stem cells (blue) from other mouse bone marrow cells (red) assayed by Grün et al. 2016. (**b**) Specter does not rely on a single set of parameters, but performs multiple runs of landmark-based clustering using different sets of landmarks of different size and different measures of similarities between cells (parameterized by *σ*). Three clusterings closely resemble the true labeling shown in (a), while one differs substantially. (**c**) Specter reconciles all individual clusterings into a consensus clustering. It clusters a carefully selected subset of cells (marked by circled stars) based on their co-association across all individual clusterings in (b), indicated by the width of the corresponding edge. The thicker an edge, the more often its two endpoints were placed in the same cluster. Here, the 4 red stars and the 2 blue stars correctly form 2 groups of cell, whose labels are finally propagated to the remaining cells using 1-nearest neighbor classification. The final clustering shown in (c) closely resembles the true clustering in (a).

## Results

### Specter is more accurate than competing methods

We compared the performance of Specter to representative scRNA-seq clustering methods SC3 (v1.10.1) (Kiselev et al. 2017), Seurat (v2.3.4) (Satija et al. 2015), dropClust (v2.1.0) (Sinha et al. 2018), RCA (v2.0) (Li et al. 2017), TSCAN (v1.24.0) (Z Ji and H Ji 2016), RaceID3 (v0.2.1) (Herman et al. 2018), CIDR (v0.1.5) (P Lin et al. 2017), RtsneKmeans (Duò et al. 2018) as well as to a geometric sketching based clustering approach (Hie, Cho, et al. 2019). SC3 and Seurat consistently demonstrated superior performance over competing methods in several clustering benchmarks (Duò et al. 2018; Tian et al. 2019) and are routinely used in scRNA-seq based cell type analyses. The graph-based Louvain clustering approach used by Seurat has an additional speed advantage over SC3, which applies a consensus clustering scheme to obtain particularly accurate clusterings. dropClust was recently proposed for the analysis of ultra-large scRNA-seq data sets and follows a strategy outlined above. It first reduces the size of the data to a maximum of 20,000 cells using random sampling. After a second sampling step based on Louvain clusters, it applies average-linkage hierarchical clustering on the sampled cells. Cluster labels are then transferred to the remaining cells using a Locality Sensitive Hashing forest (Bawa et al. 2005) for approximate nearest neighbor searches. In contrast, the geometric sketching algorithm proposed in Hie, Cho, et al. 2019 samples cells evenly across the transcriptional space rather than proportional to the abundance of cell types as uniform sampling schemes do. Experiments in Hie, Cho, et al. 2019 demonstrated that clustering a geometric sketch using the graph-based Louvain algorithm followed by propagating labels to the remaining cells via *k*-nearest-neighbor classification accelerates clustering analysis and yields more accurate results than uniform sampling strategies. We include the same geometric sketching based clustering method in our benchmark and refer to it simply as geometric sketching throughout the text. We further included methods RCA, TSCAN, RaceID3, and CIDR to cover a diverse set of algorithms commonly used to cluster scRNA-seq data (see recent benchmarks Duò et al. 2018; Tian et al. 2019; Freytag et al. 2018), from nearest neighbor based graph clustering to hierarchical clustering to *k*-medoids to model-based clustering. Finally, we included general-purpose *K*-means clustering (RtsneKmeans) as a baseline that performed surprisingly well in Duò et al. 2018 compared to methods specifically developed for clustering scRNA-seq data.

### Data sets and evaluation

We evaluated Specter and competing methods on 21 public scRNA-seq data sets and 24 simulated data sets (Supplemental Tables S1, S2). The former includes 16 data sets for which cell type labels were inferred in the original publication from clusterings of scRNA-seq measurements which typically underwent manual refinement and annotation as well as all but one real data sets that were used in (Duò et al. 2018) to benchmark clustering methods based on cell phenotypes defined independently of scRNA-seq. Identically to Duò et al. 2018, we used “true” cell types annotated by FACS sorting in the *Koh* data set, and partitioned cells by genetic perturbation and growth medium in the *Kumar* data set. In data sets *Zhengmix4eq* and *Zhengmix4uneq* the authors of Duò et al. 2018 randomly mixed equal and unequal proportions, respectively, of pre-sorted B-cells, CD14 monocytes, naive cytotoxic T cells and regulatory T cells. Data set *Zhengmix8eq* additionally contained roughly equal proportions of CD56 NK cells, memory T cells, CD4 T helper cells, and naive T cells. Again, annotated cell types were used as reference partitioning of cells in the evaluation. We excluded a single data set from (Duò et al. 2018) in which ground truth labels correspond to collection time points which all methods tested in Duò et al. 2018 failed to reconstruct. Data sets vary in size and number of cell populations and are described in Supplemental Table S1. We used Splatter (Zappia et al. 2017) to simulate 24 data sets that varied in the relative abundance of cell types that were either all equal (G*eq*), unequal (G*neq*), or based on cell type abundances among peripheral blood mononuclear cells (PBMCs) in healthy individuals (G*pbmc*), in number of cells (N*1k*, N*2k*, N*5k*), and in the probability of a gene being differentially expressed in a group, which was either 0.01 (DE*1*), 0.02 (DE*2*), 0.05 (DE*5*), or differed between groups (DE*neq*). Following Zappia et al. 2020, we set the number of genes to 1, 000 or 10, 000 (D*10k*). Supplemental Table S2 lists the characteristics of all simulated data sets.

We apply standard and uniform preprocessing (Duò et al. 2018) on all real and simulated data sets, including natural log-transformation of gene counts after adding a pseudo-count of 1, selection of top 2,000 most variable genes (omitted for simulated data sets with less than 2,000 genes), followed by dimensionality reduction to 100 principle components (Vijayan 2020). We show results for Specter when using 20 ensemble members (*Specter20E*) and 50 ensemble members (*Specter50E*), which we motivate below through experiments addressing the dependence of Specter’s accuracy on the number of ensemble members. The results for these two variants are nearly identical and we therefore simply refer to them as Specter unless we explicitly distinguish these two settings. Due to our clustering ensemble scheme, no additional tuning of parameters is required to apply Specter to the 45 data sets. The geometric sketching based Louvain clustering is provided with the same preprocessed data as Specter, all other methods are run with their built-in data preprocessing. Consistent with the original publication (Hie, Cho, et al. 2019), geometric sketches ranging from 2% to 10% of the original number of cells were computed and clustered as described above. All methods were provided the correct number of clusters or corresponding parameters were tuned accordingly. All experiments were run on a Intel Xeon CPU @2.30GHz with 320 GB memory. Methods SC3, RCA, RaceID3, and CIDR failed to run on the three largest data sets that included more than 450, 000 cells (Supplemental Table S1) due to insufficient memory. In fact, with a running time that grows cubic with the number of cells, SC3 is not designed for large data sets. On data set *chen*, for example, it takes SC3 five hours to cluster 14,000 cells. Similarly, on the three largest data sets we replaced the R implementation of the Louvain clustering algorithm called in the Seurat clustering pipeline by a more efficient python implementation of the same algorithm in the scanpy package (v1.4.6) (Wolf et al. 2018). scanpy was specifically designed for the analysis of large-scale gene expression data sets and was used originally (Cao, Spielmann, et al. 2019) to identify cell types in data set *trapnell* comprising more than 2 million cells.

Consistent with other benchmarks (see, e.g., Duò et al. 2018; Sinha et al. 2018; Freytag et al. 2018), we used the Adjusted Rand index (ARI) (Hubert and Arabie 1985) to measure the similarity between the inferred clusterings and the ground truth clustering that is based on the biological cell types annotated or pre-sorted in the original study or was provided by the simulator. We additionally applied routinely used (Freytag et al. 2018) clustering metrics Normalized Mutual Information (NMI) (Studholme et al. 1999) and a homogeneity score (Rosenberg and Hirschberg 2007) to provide a more detailed analysis of clustering performance.

### Evaluation on real data

Consistent with previous benchmarks, SC3 and Seurat overall outperform existing methods, with RCA showing a competitive performance especially with respect to homogeneity scores (Figure 2 and Supplemental Figures S1, S2). Specter, however, improves mean clustering accuracy over both methods, in all three metrics. The biggest improvement can be observed with respect to ARI and homogeneity scores, whose mean values (excluding the three largest data sets where SC3 failed to run) achieved by Specter (*Specter50E*) are 0.88 and 0.89, respectively, compared to 0.69 and 0.76 for Seurat and 0.78 and 0.84 for SC3. Overall, most methods achieved higher scores in NMI than in the other two metrics. On 17 out of 21 real data sets, Specter obtained more accurate clusterings in all three metrics than Seurat and without exception achieved higher ARI scores than sampling based methods dropClust and geometric sketching, even when sampling as many as 10% of cells in the latter approach. A similar preeminence can be observed when applying metrics NMI and homogeneity score. On many instances, the improvement was substantial. Results for smaller sketch sizes are shown in Supplemental Figure S3. In fact, on average methods dropClust and geometric sketching achieved slightly lower scores with respect to all three metrics than baseline algorithm RtsneKmeans that simply applies standard *k*-means clustering on t-SNE projected cells. Note that the ground truth labeling of cell types in data sets *trapnell, CNS*, and *saunders* was obtained in the original publication using Seurat or its underlying Louvain clustering algorithm. Despite the additional manual refinement applied in (Zeisel et al. 2018; Cao, Spielmann, et al. 2019; Saunders et al. 2018), this might positively impact the evaluation results of Seurat and the geometric sketching based Louvain clustering. On several instances, Specter achieved considerably higher ARI scores than SC3, while on others their performance was similar (within less than 10% difference in ARI). Note, however, that SC3 is not designed to cluster large data sets and had to be excluded from the comparison on the three largest data sets for computational reasons.

**Figure 2:**
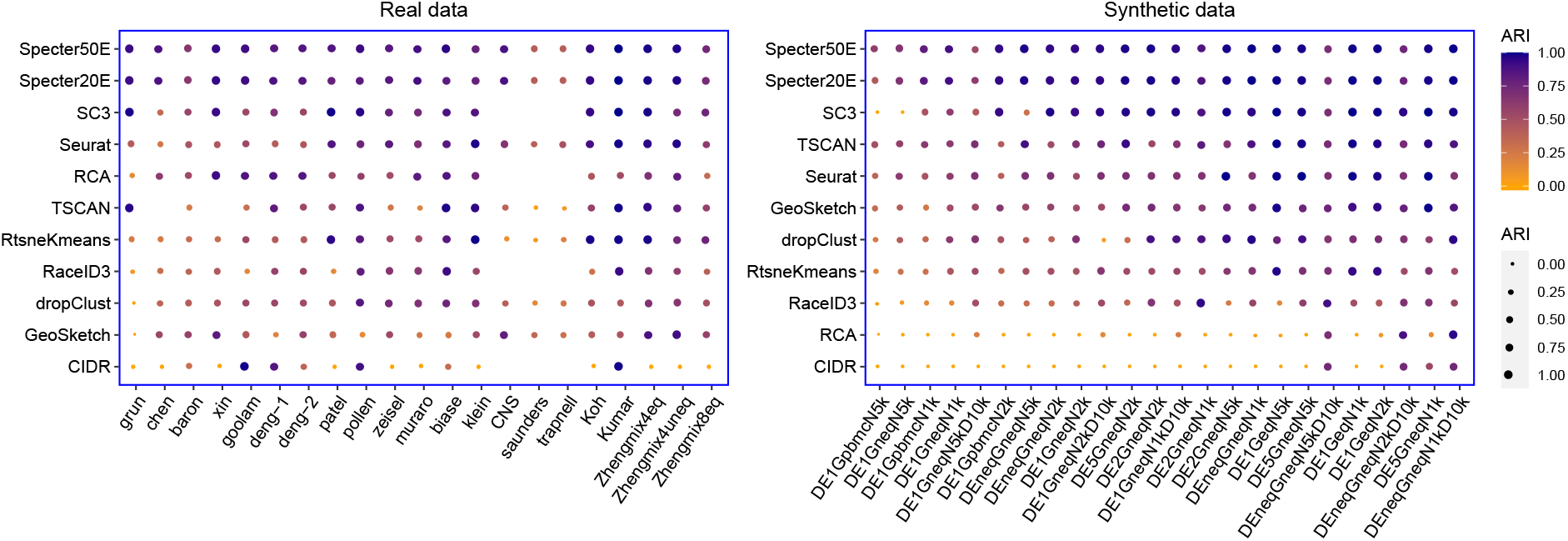
Clustering performance measured in ARI of Specter and competing methods on real and synthetic scRNA-seq data sets. Methods are ordered by mean ARI score across data sets decreasing from top to bottom. In the calculation of mean scores we excluded for each method the data sets where the method did not run successfully. For the rightmost 5 real data sets ground truth labels are based on cell phenotypes defined independently of scRNA-seq (see Supplemental Table S1). Synthetic data sets are ordered from left to right by increasing mean ARI over all methods. SC3, RCA, RaceID3, and CIDR failed to run on the three largest data sets *CNS, saunders*, and *trapnell* due to insufficient memory. TSCAN failed to run on data sets *chen* and *skin* for unknown reasons. Geometric sketching refers to the Louvain clustering of 10% of the cells sampled using geometric sketching. Results for different sketch sizes are shown in Supplemental Figures S3.

### Evaluation on simulated data

As expected, simulated data sets G*pbmc* that reflect the unbalanced cell type composition among PBMCs pose the biggest challenge to clustering algorithms, while uniform cell type abundances (G*eq*) or a larger number of marker genes (DE*neq*\*D*10k* or DE5) facilitate the detection of transcriptionally distinct groups of cells (Figure 2, Supplemental Figures S1 and S2). Consistent with results on real data sets, Specter achieved highest accuracy in terms of mean ARI, NMI, and homogeneity score across 24 simulated data sets, with scores in NMI being generally higher for most methods than in the other two metrics. Again, SC3 performs best among remaining methods in terms of mean ARI and mean homogeneity score which may be attributed to a consensus clustering scheme that it applies similarly to Specter. With respect to NMI, Seurat and TSCAN achieved slightly higher mean scores than SC3, mainly due to the two presumably most difficult instances where SC3 returned clusterings with a score of 0 (in all 3 metrics) and is thus no better than a random partition of cells. Seurat performed well on data sets with equal cell type proportions (G*eq*) and on data sets where groups are identified by a large number of marker genes (DE*5*) whereas a substantial drop in ARI and homogeneity score can be observed on the remaining data sets. Seurat’s NMI scores exhibit a similar but less pronounced pattern. Geometric sketching, which uses the same Louvain clustering algorithm as Seurat, behaves similarly. TSCAN performed better on synthetic than on real data sets (in all 3 metrics), while the opposite is true for RCA. The baseline algorithm RtsneKmeans yields remarkably accurate clusterings, especially on data sets with balanced cell type composition. On more difficult data sets, however, its accuracy drops significantly compared to several methods tailored to scRNA-seq analysis, especially in terms of ARI and homogeneity score. dropClust, on the other hand, achieved mean accuracy scores on synthetic data sets which are close to the baseline algorithm’s ones (ARI 0.63 vs 0.57, homogeneity score 0.65 vs 0.63, NMI 0.67 vs 0.71).

Finally, we illustrate in Supplemental Figure S4 how higher performance scores translate into a more meaningful representation of cell types.

### Specter facilitates robust landmark-based clustering of single cells

In addition, we compared Specter to the original implementation of the landmark-based spectral clustering (LSC) algorithm and dissect the relative contribution of our hybrid landmark selection strategy, the clustering ensemble approach and the novel selective sampling scheme (see the “Methods” section) to the overall improvement in performance by Specter (using 50 ensemble members). We show results for three variants of Specter in which we either replace the *k*-means based landmark selection or the selective sampling approach by standard random sampling, or in which we omit the clustering ensemble step altogether. Supplemental Figures S5 and S6 demonstrate the effectiveness of our adoptions and extensions of the original algorithm to the analysis of scRNA-seq data. Across all 24 simulated data sets, Specter achieved a higher ARI (mean ARI 0.89) than LSC (mean ARI 0.59) (Supplemental Figure S5). In fact, even without the benefit of a clustering ensemble, further algorithmic adjustments implemented in Specter such as a modified bandwidth of the Gaussian kernel yielded an improvement over LSC on 19 out of 24 data sets. When disabling the clustering ensemble approach in Specter, however, its performance decreased consistently, on several data sets the decrease in ARI was substantial. Similarly, on 21 out of 24 data sets the selective sampling in Specter was more effective in terms of ARI than random sampling. On two instances with unbalanced cell type compositions (*pbmc*), the score more than doubled. Remarkably, coupled with random sampling (instead of selective sampling), the consensus clustering obtained from a clustering ensemble was often even less accurate than a single clustering. The hybrid *k*-means based landmark selection led to an improvement in ARI on all but one data sets (Supplemental Figure S6). In many cases this improvement was substantial, especially on difficult instances with unbalanced cell type compositions (*pbmc*, G*neq*).

In Supplemental Figure S7 we further addressed the dependence of Specter’s accuracy on the number of ensemble members from which Specter computes a consensus clustering. Consistent with our observation in Supplemental Figure S5, the clustering ensemble approach yielded on average more accurate results on the 24 simulated data sets than relying on a single clustering for each data set. Even a small number of ensemble members (e.g. 10) improved clustering accuracy substantially, while only minor improvements were achieved when increasing their number further to more than 20 ensemble members. Nevertheless, a clustering ensemble of size 200 yielded highest mean ARI with lowest score variance.

Finally, we demonstrate robustness of Specter to the choice of parameter *γ* that controls the bandwidth of the Gaussian kernel that is set differently in Specter compared to LSC (see the “Methods” section). Even though this parameter is randomly selected from interval [0.1, 0.2] consistently across all 45 data sets in this benchmark, Supplemental Figure S8 shows that with very few exceptions choosing *γ* from different intervals would yield nearly identical results.

### Specter is sensitive to rare cell populations

In this section, we evaluate Specter’s sensitivity to rare cell populations. We devised three simulation experiments with increasing degree of difficulty. First, we repeated the experiment performed by Sinha et al. 2018 and randomly sampled a rare population of cells that comprise between 1% and 10% of total cells. More specifically, starting from two (equal size) groups of 2000 cells each that were simulated using Splatter (data set RareCellExp1 in Supplemental Table S2), we randomly downsample one group to comprise 1 — 10% of the total number of cells. We repeat the experiment five times for each group and similar to Sinha et al. 2018 report the average *F*_1_ score over the 10 runs in Supplemental Figure S9. The *F*_1_ score denotes the harmonic mean of the recall and precision, which we define identically to Sinha et al. 2018 with respect to the predicted cluster with the largest number of rare cells. While several methods performed well on a sample of 10% of cells (SC3 being a notable exception), only Specter and Seurat are able to accurately detect a cell population that is composed of only 1% of cells. Additionally, we performed an experiment in which we randomly sampled cells from a group that is initially smaller (1,000 cells) than the second group (9,000 cells) (data set RareCellExp2 in Supplemental Table S2). Compared to the previous experiment, the rare population of cells will then occupy a smaller transcriptional space relative to the larger group, which may represent a more realistic, but also a more challenging scenario for clustering methods. Note that the smaller group initially consists of 10% of total cells and was therefore downsampled to comprise 1 — 5% of cells. Again, each sampling experiment was repeated 10 times and average *F*_1_ scores are shown in Supplemental Figure S10. Here, several methods obtained an *F*_1_ score of close to 0 even when sampling 5% of cells, underlining the added difficulty of clustering unbalanced cell types. After further reducing the abundance of the rare cell type to 1%, only Specter achieved an almost perfect *F*_1_ score (0.96), followed again by Seurat with an *F*_1_ score of 0.78. In the most challenging scenario, we randomly downsampled naive cytotoxic or regulatory T cells that partly overlap in the Zhengmix4eq data set (see Supplemental Figure S11) to comprise 1%-10% of the total number of cells and repeated this experiment five times for each group. Average *F*_1_ scores are shown over the 10 runs in Supplemental Figure S12. Even though Specter consistently demonstrates highest accuracy among all methods, its *F*_1_ score monotonically decreases from close to 1 for 10%, to 0.26 for just 1% of cells, highlighting the intrinsic difficulty of detecting rare cell types that are transcriptionally similar to more abundant cell populations.

Finally, we confirmed Specter’s sensitivity to rare cell types on a rare population of inflammatory macrophages that was reported and experimentally validated by Hie, Cho, et al. 2019. In (Hie, Cho, et al. 2019), the authors applied Louvain clustering to a geometric sketch of 20,000 cells sampled from a data set of 254,941 umbilical cord blood cells. In their experiments the authors observed that this rare subtype is invisible to Louvain clustering, the algorithm used by Seurat, unless cells are initially sampled evenly across transcriptional space to better balance the abundance of common and rare cell types. In contrast, Specter reveals a similar population of inflammatory macrophages characterized by the same set of marker genes *CD74, HLA-DRA, B2M* and *JUNB* (AUROC > 0.9) without any prior preprocessing (Figure 3).

**Figure 3:**
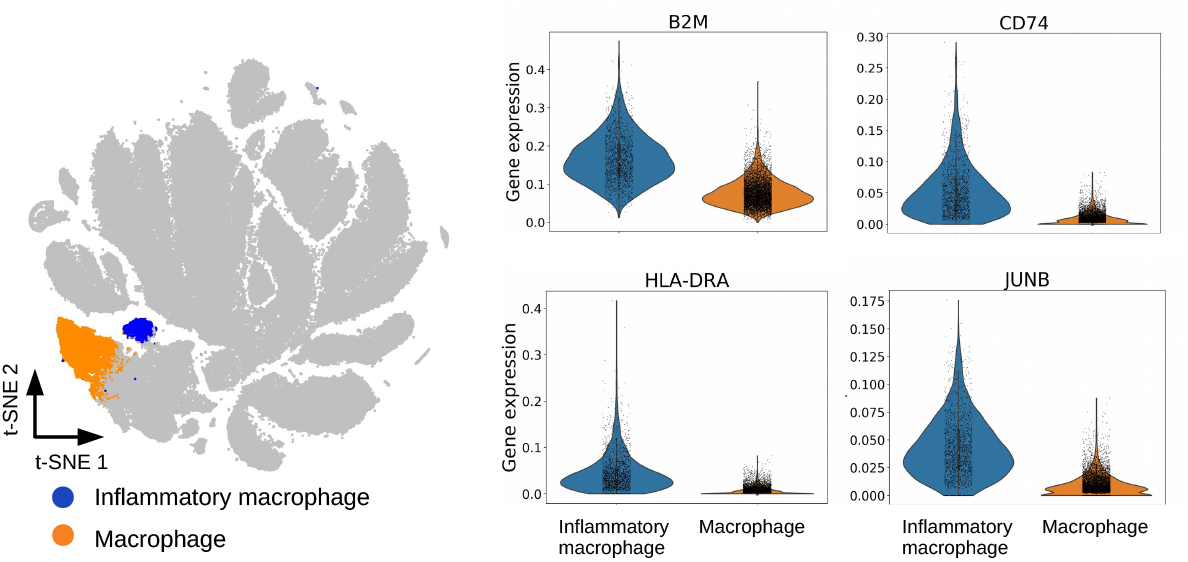
Clustering of 254,941 umbilical cord blood cell by Specter. Among macrophages defined by CD14 and CD68 marker gene expression, Specter detects a rare subpopluation of inflammatory macrophages that was recently discovered (Hie, Cho, et al. 2019) (*left*). This rare subtype can be distinguished in Specter’s clustering by the expression of the same set of inflammatory marker gene expression (*CD74, HLA-DRA, B2M*, and *JUNB*) used for its identification in Hie, Cho, et al. 2019.

### Specter utilizes multi-modal data to resolve subtle transcriptomic differences

In this Section, we demonstrate the ability of Specter to utilize complementary information provided by multi-modal data to refine the clustering of single cells. More specifically, we re-analyzed two public data sets of 4,292 healthy human peripheral blood mononuclear cells (PBMC) (Mimitou et al. 2019) and 8,617 cord blood mononuclear cells (CBMC) (Stoeckius et al. 2017), for which both mRNA and protein marker expressions (ADT, antibody-derived tags) were measured simultaneously using CITE-seq (Stoeckius et al. 2017). In these experiments, the authors used 49 and 13 antibodies, respectively, that recognize cell-surface proteins used to classify different types of immune cells.

Consistent with previous analyses of CITE-seq data (Satija 2019; Kim et al. 2020), we used the Seurat R package (Butler et al. 2018) to preprocess RNA and ADT counts. We normalized ADT expression using centered log-ratio (CLR) transformation and log-transformed RNA counts after adding a pseudocount of 1. After selecting the top 2,000 most variable genes, the expression of each gene was scaled to have mean expression 0 and variance 1, followed by dimensionality reduction to 20 principal components.

Doublets in the PBMC data set were removed using the same cell hashing-based approach with identical parameters as in Kim et al. 2020. Similar to the analysis in Stoeckius et al. 2017, a putative cluster of doublets coexpressing different RNA and protein lineage marker were removed from further analysis. On the CBMC data we relied on the doublet removal of Seurat performed in a prior analysis (Satija 2019) of this data set.

We annotate clusters based on differential expression of marker genes (Wilcoxon rank-sum test) for immune cell types listed in Supplemental Table S3. The analysis of both data sets is documented at https://github.com/canzarlab/Specter.

On both data sets, both Seurat and Specter fail to accurately distinguish naive CD4 T cells and CD8 T cells based on transcriptomic data alone (Figures 4; Supplemental Figure S13). Many CD4-/CD8+ T cells identified by protein measurements (ADT) in the CBMC data set are wrongly grouped together with CD4 T cells by Seurat and Specter. Similarly, CD4 and CD8 T cells are mixed in the PBMC data set by both methods.

**Figure 4:**
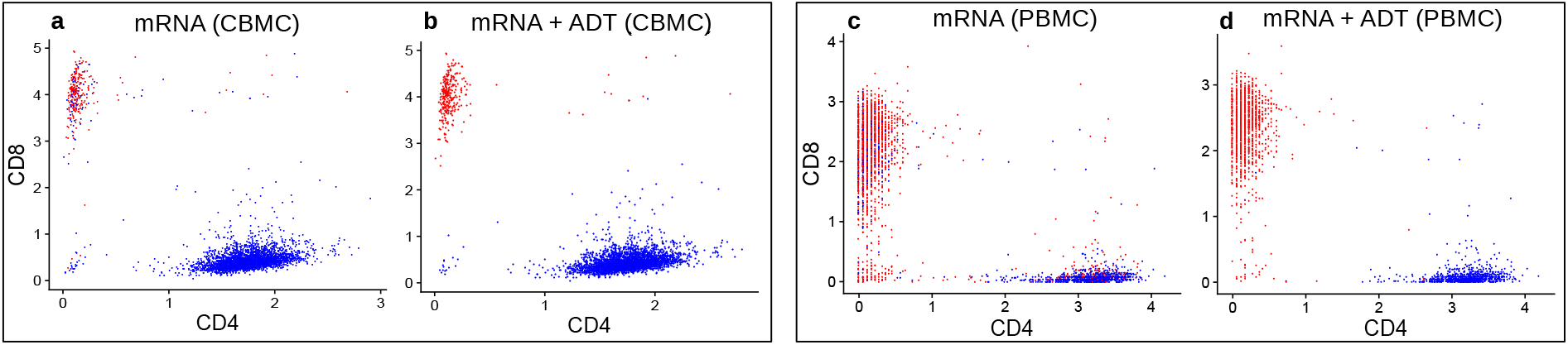
Comparison of unimodal and joint clustering by Specter. CBMCs (left box) and PBMCs (right box) with coordinates of protein expression (ADT) along CD4 and CD8 axis. Cells are clustered by Specter into CD4 T cells (blue) and CD8 T cells (red) either based on mRNA expression alone (**a, c**) or jointly from mRNA and surface protein expression (**b, d**). The mixing of CD4 T cells and CD8 T cells in the mRNA based clustering is corrected through the co-association of both modalities by Specter.

On the other hand, dendritic cells and megakaryocytes cannot be identified in the CBMC data set based on protein marker expression, see analysis using Seurat (Satija 2019). Similarly, Figure 5 shows that ADT-based clustering by Specter is not able to separate CD14+ from FCGR3A+ Monocytes nor megakaryocytes from other cell types in the PBMC data set. This can be analogously observed in the clustering by Seurat (Supplemental Figure S13).

**Figure 5:**
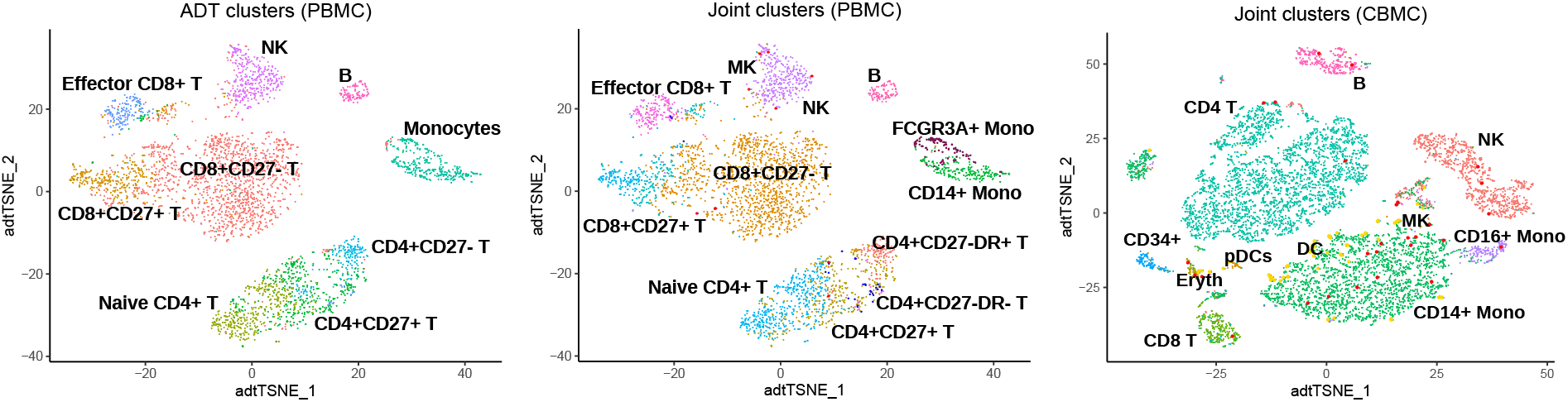
t-SNE visualization of clusters identified by Specter. Clusters of PBM cells were inferred from protein expression (ADT) alone (left) or from combined mRNA and protein expression (middle). In contrast to the joint clustering of both modalities, ADT-based clustering cannot discriminate CD14+ and FCGR3A+ Monocytes, does not detect megakaryocytes (red) and does not allow to discriminate between CD27-DR+ and CD27-DR-subpopulations of CD4+ T cells. The simultaneous clustering of RNA and protein expresssion in CBM cells (right) additionally reveals a rare population of megakaryocytes (red).

We therefore aimed to correct and improve the individual clusterings of RNA and surface marker protein measurements by combining the two distinct species through our clustering ensemble approach. In particular, Specter first produces an identical number of clusterings (here 200) for each modality. It then combines the transcriptome-based clusterings and the protein-based clusterings through a co-association approach (see the “Methods” section).

The joint clustering of RNA and protein expression by Specter profits from both modalities, yet differs from both unimodal analyses: On the PBMC data set, an ARI score of 0.78 comparing multimodal and RNA-based clustering and a score of 0.72 between multimodal and ADT-based clustering indicate complementary aspects of cellular identity utilized in their joint clustering. On the CBMC data set, higher ARI scores of 0.87 and 0.91 between the multimodal clustering and RNA and ADT-based clusterings, respectively, reflect a higher agreement between the two modalities.

More specifically, the joint clustering of RNA and protein expression of CBM and PBM cells allows Specter to more accurately separate CD4 T cells and CD8 T cells compared to a simple transcriptome-based clustering (Figure 4). In contrast to ADT expression based clustering of PBM cells, the joint clustering of RNA and surface protein expression by Specter correctly identifies megakaryocytes, CD14+, and FCGR3A+ Monocytes (Supplemental Table S3 and Figure 5). In addition, only the combined clustering of ADT and RNA allows Specter to discriminate between CD27-DR+ and CD27-DR-subpopulations of CD4+ memory T cells. In contrast to the clustering of protein data of CBM cells, Specter also correctly detects dendritic cells and megakaryocytes based on the markers listed in Supplemental Table S3 (see Figure 5).

We compare the joint clustering by Specter to the results of CiteFuse (v0.99.10) (Kim et al. 2020), a method that was recently proposed specifically for the computational analysis of single cell multimodal profiling data. As proposed initially for the combination of (bulk) genome-wide measurements across, e.g., patients (Wang et al. 2014), CiteFuse applies the similarity network fusion algorithm to combine RNA and ADT expression of single cells and then clusters the fused similarity matrix using spectral clustering. We ran CiteFuse as originally described in (Kim et al. 2020) including the removal of doublets and the (internal) selection of highly variable genes.

Overall, the clusters of CBM and PBM cells as computed by Specter and CiteFuse are highly similar, as indicated by a high ARI score of 0.94 and 0.86 for the two data sets (Supplemental Figures S14 and S15). In both data sets, however, only Specter is able to identify a rare population of megakaryocytes (Supplemental Table S3). Furthermore, in contrast to the analysis performed in Kim et al. 2020, CiteFuse was not able to discriminate beween CD27-DR+ and CD27-DR-subpopulations of CD4+ memory T cells in the PBMC data set, neither when using identical parameters as in Kim et al. 2020 nor when applying more conservative parameters in the doublet removal (parameters taken from CiteFuse tutorial (Y Lin and Kim 2020)) (Supplemental Figure S16). The authors of Kim et al. 2020 attribute this discrepancy to a different selection of highly variable genes applied in an earlier version of the software used to produce the results in Kim et al. 2020.

The major advantage of Specter over CiteFuse, however, is its speed and scalability. CiteFuse requires 15 minutes and nearly 2 hours to jointly cluster the 3,880 PBM cells and 7,895 CBM cells (after doublet removal), respectively, and is thus not expected to scale well on much larger data sets due to the computational expensive fusion of networks. In contrast, Specter returns a high resolution clustering of the two data sets in just 20 and 50 seconds, respectively.

### Scalability

Here, we demonstrate the scalability of Specter to large single-cell data sets. To experimentally confirm the theoretical linear-time complexity of our algorithm, we devised different size simulated data set containing between 1,000 and 1 million cells (with characteristics DE1Geq, see Supplemental Table S2.). As expected (Cai and Chen 2011), the landmark-based sparse representation of the data allows to compute a spectral embedding in linear time (see Supplemental Figure S17). Furthermore, the experiment confirms that our novel selective sampling strategy reduces the quadratic complexity of the hierarchical clustering step that reconciles multiple ensemble members (see the “Methods” section) to an overall linear dependence on the number of cells. As expected, the rate of increase in running time, i.e. the slope of the lines shown in Supplemental Figure S17, is larger when Specter includes multiple clusterings (here 20) in the ensemble scheme. More precisely, we observed a linear increase in running time with the size of the clustering ensemble, that is, with the number of independent runs of the core algorithm (Supplemental Figure S18). However, as shown in our experiments assessing the importance of individual algorithmic components in Specter, a relatively small number of runs is sufficient to improve accuracy of the resulting consensus clustering substantially. Even more, the independent computation of individual clusterings in an ensemble lend themselves to parallel processing. In Supplemental Figure S19 we therefore explored how the use of multiple threads can speed-up the clustering ensemble approach and thus counterbalance the inclusion of an increasing number of ensemble members. With just 4 threads, the time required to compute a consensus clustering from 50 individual clusterings of 100, 000 cells reduced from around 92 seconds to just 34 seconds. Increasing the number of threads further has a decreasing effect on total running time, reaching 15 seconds total computation time using 20 threads. Again, we observed a roughly linear increase in running time with increasing sample size for fixed number of threads (Supplemental Figure S20), where 4 threads reduced the running time of 50 runs in the clustering ensemble to a time that is nearly identical to the time a single threads needs to compute a consensus clustering from 20 ensemble members.

In Figure 6 we compared Specter’s running time to all methods that ran successfully on the three largest real data sets. For all methods except TSCAN and dropClust we measured the running time of the core algorithm and exclude preprocessing. The time Specter required to preprocess the data (using a single thread), including log-transformation, the selection of highly variable genes, and principle component analysis, is negligible (Supplemental Tables S4 and S5). Seurat was run with a call to the more efficient scanpy implementation of the Louvain clustering algorithm. Even in single-threaded mode, Specter’s running time that included 20 individual clusterings of 1 million cells is with 7.6 min considerably faster than Seurat which required 23 min for a single Louvain-based clustering of the same set of cells (Supplemental Table S4). Note that 20 ensemble members were used by Specter in Figure 2 (and Supplemental Figures S1, S2) to achieve overall more accurate clusterings than competing methods. With just 4 threads Specter’s running time further drops to 3.2 min (Supplemental Figure S20), whereas Seurat’s clustering algorithm cannot be run with multiple threads. dropClust required 6.8 minutes to preprocess and cluster 1 million cells, but is not able to make use of multiple threads. The running time of geometric sketching increases the fastest while RtsneKmeans is as expected the slowest method (Supplemental Figure S21). As one potential use of clustering results, visualization by FIt-SNE (Linderman et al. 2019) required around 8 min for 1 million cells (Supplemental Table S5).

**Figure 6:**
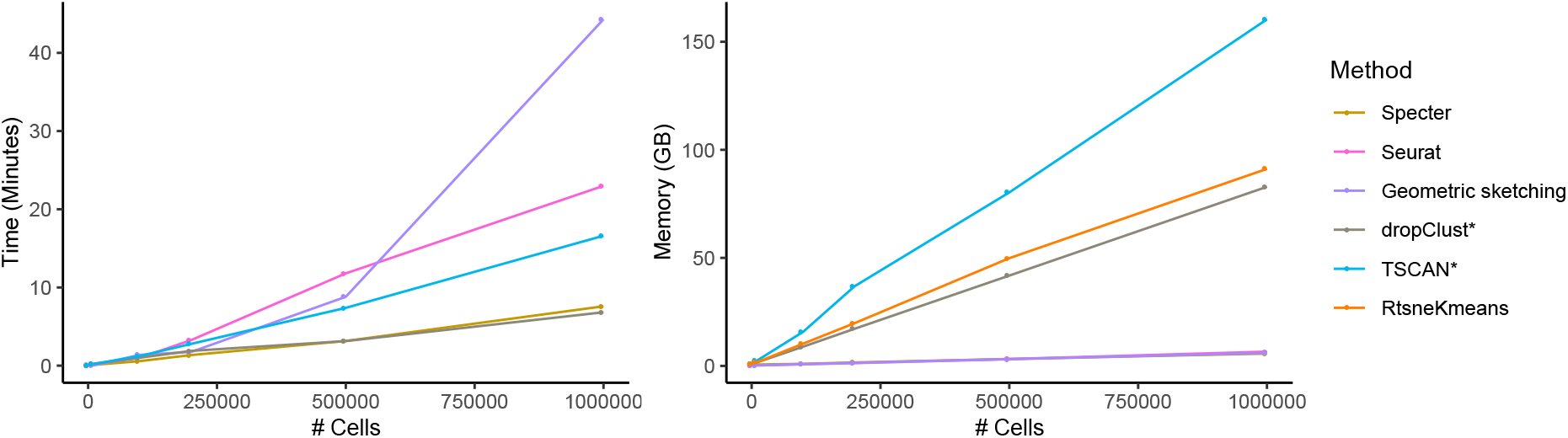
Runtime and peak memory usage as a function of sample size. Seurat was run with a call to the more efficient SCANPY implementation of the Louvain clustering algorithm. Running times exclude preprocessing for all methods except TSCAN and dropClust, whose implementation did not allow to isolate the core algorithm. Memory usage of Specter, Seurat, and geometric sketching are nearly identical and cannot be distinguished in this plot. For ease of visualization we show runtime results of method RtsneKmeans in Supplemental Figure S21.

Finally, Supplemental Table S6 gives the CPU times in minutes on the three largest real data sets used in this study. Again, we excluded preprocessing for all methods except TSCAN and dropClust. We additionally report the total running time of Specter including all prior preprocessing. In this analysis of real data sets, we exploited the full performance potential of Specter and used 20 threads to compute consensus clusterings from 50 individual runs, which outperformed all other methods in terms of accuracy in Figure 2 and Supplemental Figures S1, S2. In this setting, Specter required around 15 min to cluster 2 million cells (23 min including single-threaded preprocessing) and was 5-10 times faster than Seurat that is not able to utilize multiple threads. On the largest data set, dropClust was with just 12 minutes of total computation time using just a single thread the fastest method. In contrast to Specter, however, dropClust considers only around 1% of the data (20,000 cells) and its simplified model comes at the cost of a substantial loss in accuracy (see Figure 2 and Supplemental Figures S1, S2). Again, RtsneKmeans is the slowest among methods that terminate successfully on these large data sets.

Furthermore, Figure 6 shows peak memory usage as a function of number of cells on the same simulated data sets used to evaluate runtime performance. Together with Seurat and geometric sketching, Specter required the least amount of memory (less than 7 GB for 1 million cells), while memory usage of methods TSCAN and dropClust increased rapidly for data sets containing more than 200,000 cells.

## Discussion

We have introduced Specter, a novel method that identifies transcriptionally distinct sets of cells with substantially higher accuracy than existing methods. We adopt and extend algorithmic innovations from spectral clustering, to make this powerful methodology accessible to the analysis of modern single-cell RNA-seq data sets. We have demonstrated the superior performance of Specter across a comprehensive set of public and simulated scRNA-seq data sets and illustrated that an overall higher accuracy also implicates an increased sensitivity towards rare cell types. At the same time, its linear time complexity and practical efficiency makes Specter particularly well-suited for the analysis of large scRNA-seq data sets. Besides technological advances, the integration of cells from multiple experiments spanning different tissues or diseases may yield data sets with massive numbers of cells. Coupled with data integration methods such as Scanorama (Hie, Bryson, et al. 2019) or Harmony (Korsunsky et al. 2019) that can remove, e.g., tissue-specific differences, Specter can help to leverage such reference data sets to reveal hidden cell types or states. When combining different samples from the same experiment, simpler linear methods such as ComBat (Johnson et al. 2006) might be preferable (Luecken and Theis 2019) to correct for batch effects between samples prior to identifying groups of cell with distinct gene expression profiles using Specter.

Furthermore, we have illustrated how the flexibility of its underlying optimization model allows Specter to harness multimodal omics measurements of single cells to resolve subtle transcriptomic differences between subpopulations of cells. The application of our cluster ensemble scheme to the joint analysis of multimodal CITE-seq data sets yielded a slightly more fine-grained distinction of cell (sub-)populations compared to the recently proposed multimodal clustering method CiteFuse. More importantly, in contrast to CiteFuse whose running time increased ≈ 8 fold after doubling the number of cells, Specter will scale well to much larger data sets produced by droplet-based approaches that can measure multiple modalities of up to millions of cells together. While the consensus clustering approach applied by Specter can in principle integrate the ensemble of clusterings generated from various molecular features, this work has focused on the combination of mRNA and protein marker expression as measured by CITE-seq or REAP-seq (Peterson et al. 2017). The practical suitability and potential limitations as well as necessary refinements of this strategy when applied to other assays that simultaneously measure, for example, accessible chromatin and gene expression (Cao, Cusanovich, et al. 2018), or more than two modalities at the same time (Clark et al. 2018), will need to be addressed in future experiments. Taken together, we believe that Specter will be useful in transforming massive amounts of (multiple) measurements of molecular information in individual cells to a better understanding of cellular identity and function in health and disease.

## Methods

Spectral clustering uses eigenvectors of a matrix derived from the distance between points (here cells) as a low-dimensional representation of the original data, which it then partitions using a method such as *k*-means. More precisely, given *n* data points 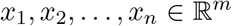 and a similarity matrix (affinity matrix) *W* = (*w_ij_*)_*n×n*_, where *w_ij_* measures the similarity between points *x_i_* and *X_j_*, the graph Laplacian is defined as *L* = *D* — *W* or *L* = *I* — *D*^-1/2^*WD*^-1/2^ in case of a (symmetric) normalized Laplacian. Here, *D* is a diagonal matrix whose entries are column sums (equivalently row sums) of *W*. Spectral clustering then uses the top *k* eigenvectors of *L* to partition the data into *k* clusters using the *k*-means algorithm.

In the following description of our algorithm we assume a given number of clusters *k*. In Specter we determine the number of clusters based on the Silhouette index (Rousseeuw 1987), which performed particularly well in recent benchmark studies (Arbelaitz et al. 2013; Chouikhi et al. 2015).

### Landmark-based spectral clustering of single cells

Several methods have been proposed to accelerate the spectral clustering algorithm (Fowlkes et al. 2004; Shinnou and Sasaki 2008; Cai and Chen 2011). In particular, Landmark-based Spectral Clustering (LSC) has been shown to perform well in terms of efficiency and effectiveness compared to state-of-the-art methods across a large number of data sets (Cai and Chen 2011). In short, LSC picks a small set of *p* representative data points 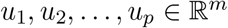, i.e. the landmarks, which it then uses to create a representation matrix 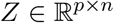 whose columns represent the original data with respect to the landmarks according to *X* ≈ *UZ*. Here, columns *i* of 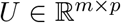 contains landmarks *u_i_* and columns *i* of *X* the original input points *x_i_*. Let the Gaussian kernel *K*(*x,y*) = exp(—||*x* — *y*||^2^/2σ^2^) measure the similarity between two points *x* and *y*, then matrix *Z* = (*z_ji_*)_*p×n*_ is computed using Nadaraya-Watson kernel regression (Härdle 1990) as

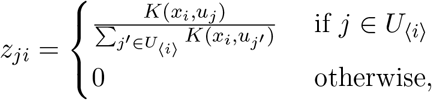

where *U*_〈*i*〉_ is the set of *r* nearest landmarks of *x_i_*. That is, *Z_ji_* is set to zero if *U_j_* is not among the *r* nearest neighbors of *x_i_*, which naturally leads to a sparse representation of the data. Motivated by non-negative matrix factorization that uses *k* (i.e. number of clusters) basis vectors to represent each data point (Xu et al. 2003), we set r to be equal to *k* in Specter (and in all experiments). Then each original point *x_i_* can be approximated by

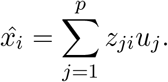

From this landmark-based representation of the *complete* data it computes the Laplacian matrix 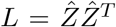, where 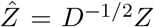 and *D* is the diagonal matrix whose (*i,i*)-entry equals the sum of the *i*th row of *Z*. Then, this graph Laplacian *L* admits a fast eigendecomposion in time *O*(*n*) as oppose to *O*(*n*^3^) in the general case, which is described in more detail in (Cai and Chen 2011).

Here, we tailor the idea of landmark-based spectral clustering to the characteristics and scale of modern scRNAseq data sets. In particular, the choice of bandwidth *σ* used in the (Gaussian) kernel to smooth the measure of similarity between pairs of data points heavily depends on the type of data and can have a strong impact on the final clustering. In the original approach, parameter *σ* is set to the average Euclidean distance between data points and their *K*-nearest landmarks, i.e. to the average value of all elements in matrix *Z*. We empirically find that replacing the average by the maximum value, i.e. by setting *σ* = *γ*×mean(max(*Z*)), where max(*Z*) denotes a vector of maximum values for each row in *Z* and *γ* a randomly chosen parameter between 0 and 1, is able to better capture the transcriptional similarity between single cells and yields more accurate clusterings of cells.

Furthermore, we pair the theoretical reduction in time complexity from 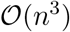 to 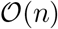 with a practical speed-up of the LSC algorithm by applying a hybrid strategy when selecting the landmarks. The choice of representative data points, here single cells, plays a crucial role in the quality of the final clustering. Random selection or *k*-means clustering were originally proposed as procedures for picking landmarks (Cai and Chen 2011). Random selection of representative cells is very efficient but often yields random sets of cells that do not represent the full data well and thus lead to poor clustering results. *k*-means, on the other hand, better takes into account the structure of the data when selecting landmark cells but its higher computational cost makes it impractical for large scRNA-seq data sets where it accounts for around 90% of the overall running time in our experiments. Our hybrid strategy seeks to balance the efficiency of random sampling and the accuracy of *k*-means based landmark selection. It first picks a set of *p*′ candidate landmarks uniformly at random with *p*′ ≪ *n* (by default, *p*′ = 10*p*), from which it subsequently selects *p* < *p*′ final landmark cells using the *k*-means algorithm. Note that despite the initial random sampling, the *full* data are represented by the final set of landmarks.

Finally, for data sets that contain a small number of clusters, we adjust the spectral embedding based on which the original data is clustered using *k*-means in the last step of spectral clustering. For a small number of clusters (e.g. *k* ≤ 4), the top *k* eigenvectors used in the original approach typically do not contain enough information to represent the full data well. In this case, we therefore use the top *k* + 2 eigenvectors to compute the spectral embedding.

### Clustering ensembles across parameters and modalities

Different data types require a different choice of parameter values and there is no general rule how to select the best one. To address this issue, we employ consensus clustering, also known in literature as cluster ensembles (Strehl and Ghosh 2003), in the same way as ensemble learning is used in supervised learning. In particular, we generate a series of component clusterings by varying the number of selected landmarks *p* and the kernel bandwidth. We randomly select parameter *γ* which controls the bandwidth of the Gaussian kernel from interval [0.1, 0.2] and choose *p* from interval

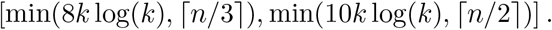

This choice of *p* is motivated by a result by Tremblay et al. 2016 who used sampling theory of bandlimited graph-signal developed in Puy et al. 2016 to prove that clustering a random subset of size *O*(*k* log(*k*)) is sufficient to accurately infer the cluster labels of all elements. To avoid sampling too many landmarks for small data sets (i.e. small number of cells *n*), we additionally set upper bounds ⌈*n*/3⌉ and ⌈*n*/2⌉ for the left and right boundaries of the interval, respectively. All clusterings produced by the different runs of our tailored LSC algorithm are then summarized in a co-association matrix *H* (Fred and Jain 2005) in which entry (*i, j*) counts the number of runs that placed cells *i* and *j* in the same cluster. We compute the final clustering through a hierarchical clustering of matrix *H*. Our LSC-based consensus clustering approach is summarized in Algorithm 1.

Different parameter choices (e.g. kernel bandwidths) provide different interpretations of the same data. In the same way as clustering ensembles can help unifying these different views on a single modality, they can help reconcile the measurements of multiple modalities, such as transcriptome and proteome, of the same cell. More specifically, Specter produces an identical number of clusterings for each modality in step 2 of Algorithm 1 which it then combines through the same co-association approach (steps 3 and 4).

#### Algorithm 1

LSC ensemble

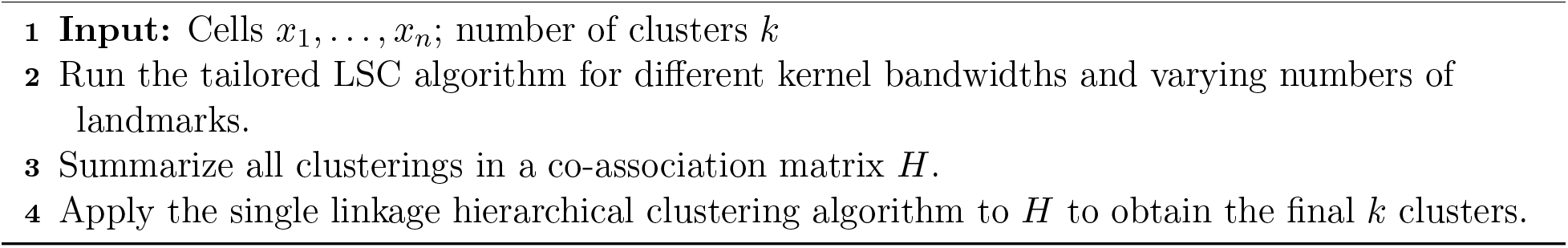

#### Time complexity

The time complexity of the tailored LSC algorithm is *O*(*n*), and single linkage hierarchical clustering requires *O*(*n*^2^) time, yielding an overall complexity of *O*(*n*^2^) for Algorithm 1, assuming *k* is small enough to be considered a constant.

### Selective sampling-based clustering ensemble

With a running time that scales quadratically with the number of cells, the application of Algorithm 1 to large-scale scRNA-seq data sets becomes infeasible. We therefore apply step 3 of our clustering ensemble approach (Algorithm 1) to a carefully selected sketch of the data. Note, however, that the co-association matrix *H* built in step 3 of the algorithm is based on cluster labels that were learned from the *full* data in step 2 using our tailored LSC algorithm. In addition, we propose a simple sampling technique that uses all clusterings computed in step 2 to guide the selection of cells.

#### Selective Sampling

Sampling cells uniformly at random is naturally fast, since the decision to include a given cell into a sketch does not depend on any other cell. At the same time, these independent decisions ignore the global structure of the data such as the abundance of different cell types and may thus lead to a loss of rare cell types (Hie, Cho, et al. 2019). We therefore propose a sampling approach that utilizes the clusterings of the data computed in step 2 of Algorithm 1 to inform the (fast) selection of cells. More specifically, Let Π = {*π*_1_,*π*_2_,...,*π*_m_}, where *π_i_* = (*π*_*i*1_, *π*_*i*2_,..., *π*_*ik*_) is the ith clustering returned in step 2 of Algorithm 1, *i* = 1,2,..., *m*. We select a sketch *S* of size 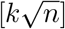 that contains roughly the same number of cells in each cluster *π_ij_*, for all *i* and *j*. This selective sampling procedure iterates through all clusters contained in all clusterings from which it randomly picks a cell not already contained in the sketch, until the size of the sketch reaches 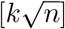 (see Algorithm 2).

##### Algorithm 2

Selective sampling

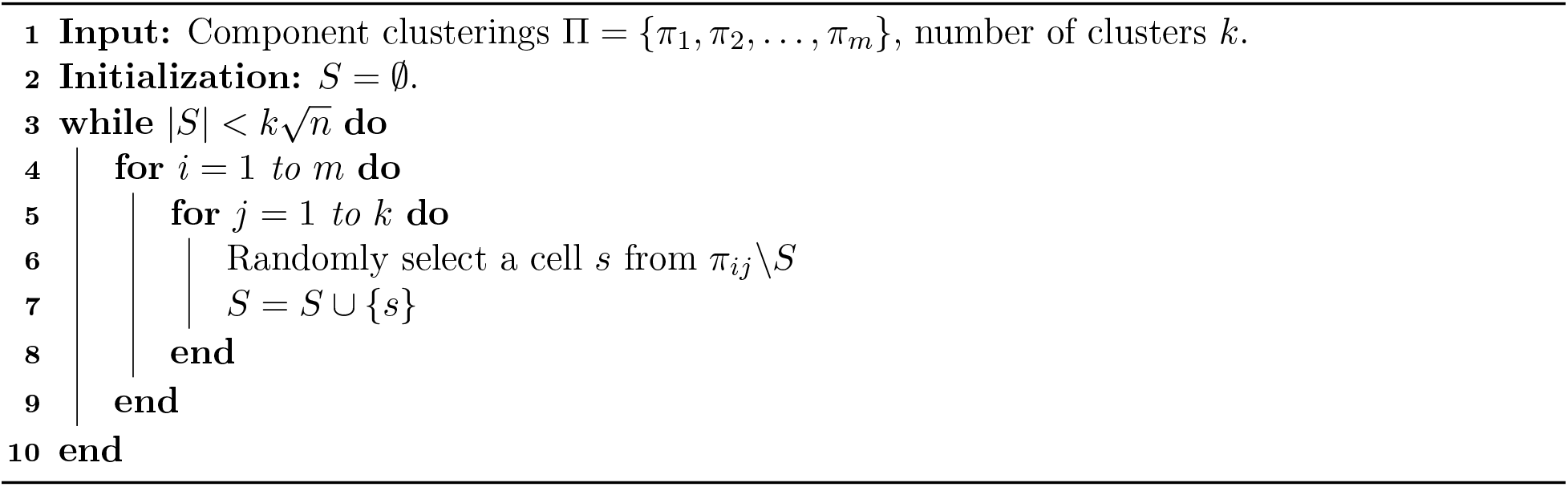

#### Inference

Given a selectively sampled sketch *S*, we apply steps 3 and 4 in Algorithm 1 to cells in *S*, using labels obtained from the full data in step 2. That is, we construct a co-association matrix whose entries count the number of times the two corresponding cells in *S* were placed in the same cluster by a run of the LSC algorithm in step 2. From this matrix, we compute a consensus clustering of *S* using hierarchical clustering and finally transfer cluster labels to the remaining cells using supervised *k*-nearest neighbors classification. That is, we assign each cell not in S to the cluster that the majority of its *k* nearest neighbors were placed in by the preceding consensus clustering of *S*. Our selective sampling-based cluster ensemble approach is summarized in Algorithm 3.

##### Algorithm 3

Selective sampling-based clustering ensemble

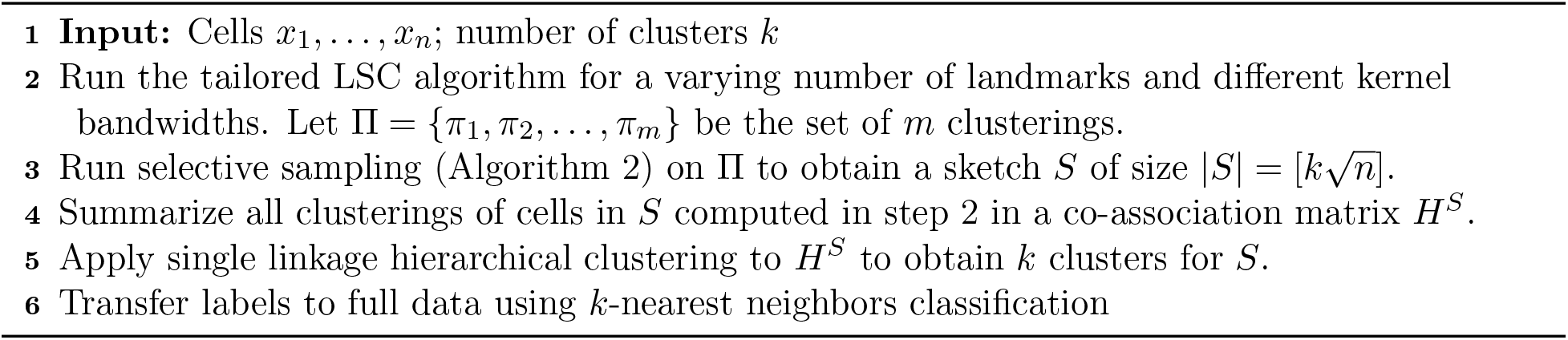

#### Time complexity

Landmark-based spectral clustering performed in step 2 of Algorithm 3 takes *O*(*n*), see above. Since we selectively sample a sketch of size 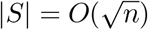 in step 3, the complexity of steps 4 and 5 now reduces to *O*(*n*). Together with the *k*-NN classification that runs in *O*(*n*) in step 6, our selective sampling based cluster ensemble scheme scales linearly with the number of cells *n*.

### Publicly available data used in this study

The original publication of data sets used in this study to assess the accuracy of Specter in comparison to existing methods are listed in Supplemental Table 1. The real data sets in Duò et al. 2018 were downloaded from https://github.com/markrobinsonuzh/scRNAseq_clustering_comparison. All other real data sets smaller than 15,000 cells were downloaded from https://hemberg-lab.github.io/scRNA.seq.datasets, the 3 largest data sets from http://mousebrain.org (*CNS*), http://dropviz.org (*saunders*), and https://oncoscape.v3.sttrcancer.org/atlas.gs.washington.edu.mouse.rna/downloads (*trapnell*). The umbilical cord blood cell data (Hie, Cho, et al. 2019) were downloaded from http://cb.csail.mit.edu/cb/geosketch.

## Software availability

The Specter software is available at https://github.com/canzarlab/Specter The Specter repository also includes all code necessary to reproduce the results of this manuscript as well as a step-by-step documentation of the analysis of the PBMC and CBMC CITE-seq data sets (Stoeckius et al. 2017; Mimitou et al. 2019) as described in this study.

## Acknowledgments

We thank Khaled Elbassioni for helpful discussion on the algorithm and Hani Jieun Kim for comments on CiteFuse results.

