## Supplemental Tables and Figures for "Linear-time cluster ensembles of large-scale single-cell RNA-seq and multimodal data"

### 1 Supplemental Tables

Supplemental Table S1: Overview of the real data sets used in this study. Names listed in the left-most column are used throughout the text. A line separates data sets in which cell type labels were inferred from scRNA-seq measurements from data set were labels are based on cell phenotypes defined independently of scRNA-seq.

| Data set | # Cells | # Populations | Description | Reference |
|---|---|---|---|---|
| *grun* | 1502 | 2 | mouse stem cells | Grün et al. 2016 |
| *xin* | 1600 | 8 | human islet cells | Xin et al. 2016 |
| *baron* | 1886 | 13 | human and mouse pancreas | Baron et al. 2016 |
| *biase* | 56 | 4 | mouse embryo devel | Biase et al. 2014 |
| *deng-1* | 268 | 6 | mouse embryo devel (RPKMs) | Deng et al. 2014 |
| *deng-2* | 268 | 6 | mouse embryo devel (Reads) | Deng et al. 2014 |
| *goolam* | 114 | 5 | mouse embryo | Goolam et al. 2016 |
| *muraro* | 2126 | 10 | human pancreas | Muraro et al. 2016 |
| *patel* | 430 | 5 | human glioblastoma | Patel et al. 2014 |
| *pollen* | 301 | 11 | human developing cortex | Pollen et al. 2014 |
| *klein* | 2717 | 4 | mouse embryo stem cells | Klein et al. 2015 |
| *zeisel* | 3005 | 9 | mouse cortex and hippocampus | Zeisel et al. 2015 |
| *chen* | 14,437 | 45 | mouse brain | Chen et al. 2017 |
| *CNS* | 465,281 | 7 | mouse central nervous system | Zeisel et al. 2018 |
| *saunders* | 665,858 | 11 | adult mouse brain | Saunders et al. 2018 |
| *trapnell* | 2,058,652 | 38 | mouse organogenesis cell atlas | Cao et al. 2019 |
| *Koh* | 531 | 9 | human embryonic stem cells | Koh et al. 2016 |
| *Kumar* | 246 | 3 | mouse embryonic stem cells | Kumar et al. 2014 |
| *Zhengmix4eq* | 3,994 | 4 | mixture of purified PBMCs | Zheng et al. 2017 |
| *Zhengmix4uneq* | 6,498 | 4 | mixture of purified PBMCs | Zheng et al. 2017 |
| *Zhengmix8eq* | 3,994 | 8 | mixture of purified PBMCs | Zheng et al. 2017 |

Supplemental Table S2: Overview of the simulated data sets used in this study. Names listed in the left-most column are used throughout the text. Data sets were simulated using Splatter (Zappia et al. 2017) and vary in number of cells (#Cells), number of genes (#Genes), number of clusters (k), the probability with which a given gene is differentially expressed in one of the cell types (marker genes), and the relative abundance of cell types that were either equal, unequal, or based on cell type abundances among peripheral blood mononuclear cells (PBMCs) in healthy individuals.

| Name | # Cells (N) | # Genes (D) | k | Probabilities of gene DE | Relative abundances (G) |
|---|---|---|---|---|---|
| DE1GeqN1k | 1,000 | 1,000 | 5 | (0.01, 0.01 0.01, 0.01, 0.01) | (0.2, 0.2, 0.2, 0.2, 0.2) |
| DE1GeqN2k | 2,000 | 1,000 | 5 | | |
| DE1GeqN5k | 5,000 | 1,000 | 5 | | |
| DEneqGneqN1k | 1,000 | 1,000 | 5 | (0.01, 0.01 0.02, 0.02, 0.05) | (0.01, 0.05, 0.14, 0.3, 0.5) |
| DEneqGneqN2k | 2,000 | 1,000 | 5 | | |
| DEneqGneqN5k | 5,000 | 1,000 | 5 | | |
| DEneqGneqN1kD10k | 1,000 | 10,000 | 5 | (0.01, 0.01 0.02, 0.02, 0.05) | (0.01, 0.05, 0.14, 0.3, 0.5) |
| DEneqGneqN2kD10k | 2,000 | 10,000 | 5 | | |
| DEneqGneqN5kD10k | 5,000 | 10,000 | 5 | | |
| DE1GneqN1k | 1,000 | 1,000 | 5 | (0.01, 0.01 0.01, 0.01, 0.01) | (0.01, 0.05, 0.14, 0.3, 0.5) |
| DE1GneqN2k | 2,000 | 1,000 | 5 | | |
| DE1GneqN5k | 5,000 | 1,000 | 5 | | |
| DE1GneqN1kD10k | 1,000 | 10,000 | 5 | (0.01, 0.01 0.01, 0.01, 0.01) | (0.01, 0.05, 0.14, 0.3, 0.5) |
| DE1GneqN2kD10k | 2,000 | 10,000 | 5 | | |
| DE1GneqN5kD10k | 5,000 | 10,000 | 5 | | |
| DE2GneqN1k | 1,000 | 1,000 | 5 | (0.02, 0.02 0.02, 0.02, 0.02) | (0.01, 0.05, 0.14, 0.3, 0.5) |
| DE2GneqN2k | 2,000 | 1,000 | 5 | | |
| DE2GneqN5k | 5,000 | 1,000 | 5 | | |
| DE5GneqN1k | 1,000 | 1,000 | 5 | (0.05, 0.05 0.05, 0.05, 0.05) | (0.01, 0.05, 0.14, 0.3, 0.5) |
| DE5GneqN2k | 2,000 | 1,000 | 5 | | |
| DE5GneqN5k | 5,000 | 1,000 | 5 | | |
| DE1GpbmcN1k | 1,000 | 1,000 | 5 | (0.01, 0.01 0.01, 0.01, 0.01) | PBMCs: DC: 0.02, NK: 0.2, B: 0.1 Mono: 0.08, T: 0.6 |
| DE1GpbmcN2k | 2,000 | 1,000 | 5 | | |
| DE1GpbmcN5k | 5,000 | 1,000 | 5 | | |
| RareCellExp1 | 4,000 | 1,000 | 2 | (0.01, 0.01) | (0.5, 0.5) |
| RareCellExp2 | 10,000 | 1,000 | 2 | (0.01, 0.01) | (0.9, 0.1) |

Supplemental Table S3: Markers used in the annotation of clusters in the CBMC and PBMC data sets. P-values indicate significance of differential expression according to a Wilcoxon rank-sum test between clusters inferred by Specter from the joint analysis of mRNA and surface protein expression.

| Cell-type | Data set | Markers |
|---|---|---|
| CD8+CD27- | PBMC | *CD8A* (p = 3.1e-15), *CD8B* (p = 4.3e-6), low *CD27* ADT |
| CD8+CD27+ | PBMC | *CD8B* (p = 3.2e-4), high *CD27* ADT |
| Naive CD4+ T | PBMC | *SELL* (Haining et al. 2008) (p = 2.6e-9) |
| CD4+CD27+ | PBMC | *IL7R* (Colpitts et al. 2009) (p = 9.4e-11), high *CD27* ADT |
| CD4+CD27-DR+ | PBMC | *IL7R* (Colpitts et al. 2009) (p = 4.4e-7), *NKG7* (Fonseka et al. 2018)(p = 1.2e-3), *GZMA* (Fonseka et al. 2018) (p = 2.0e-4) |
| CD4+CD27-DR- | PBMC | *IL7R* (Colpitts et al. 2009) (p = 1.4e-6), low expression of *NKG7* and *GZMA*; low *CD27* ADT. |
| CD14+ Mono | PBMC | *LYZ* (p = 7.5e-34), *CST3* (p = 1.4e-32) |
| FCGR3A+ Mono | PBMC | *FCGR3A* (p = 1.0e-9) |
| Megakaryocytes | PBMC | *PF4* (Lambert, Meng, Xiao, et al. 2016) (p = 1.2e-3) |
| NK | PBMC | *GNLY* (Ogawa et al. 2003) (p = 7.7e-21), *NKG7* (Turman et al. 1993)(p = 1.1e-15) |
| Dendritic cells | CBMC | *CST3* (Hruz et al. 2008) ($p$ = 4.7e-29), *CD1C* (Collin et al. 2013; Merad et al. 2013) ($p$ = 1.1e-27), and *FCER1A* (Hruz et al. 2008) ($p$ = 1.3e-27) |
| Megakaryocytes | CBMC | *PF4* (Lambert, Meng, Harper, et al. 2014) ($p$ = 1.6e-25), *PPBP* (Sakurai et al. 2016) ($p$ = 5.8e-24) |

Supplemental Table S4: Comparison of running times in minutes on simulated data. Data sets of different size were simulated using Splatter. *Running times exclude preprocessing for all methods except TSCAN and dropClust, whose implementation did not allow to isolate the core algorithm. Specter used 20 ensemble members and was run with a single thread (as all other methods). The last column (Specter+Pre) shows the total running time of Specter and all its preprocessing steps, including log-transformation, selection of highly variable genes (500), and PCA.

| #Cells | Specter | Seurat | dropClust* | Geosketch | RtsneKmeans | TSCAN* | Specter+Pre |
|---|---|---|---|---|---|---|---|
| 1k | 0.02 | 0.04 | 0.04 | 0.10 | 0.14 | 0.06 | 0.02 |
| 10k | 0.1 | 0.15 | 0.24 | 0.02 | 0.88 | 0.20 | 0.1 |
| 100k | 0.58 | 1.00 | 1.01 | 1.38 | 17.61 | 1.23 | 0.61 |
| 200k | 1.36 | 3.27 | 1.89 | 1.75 | 49.31 | 2.79 | 1.40 |
| 500k | 3.15 | 11.80 | 3.14 | 8.81 | 139.69 | 7.39 | 3.25 |
| 1m | 7.59 | 23.00 | 6.83 | 44.29 | 655.95 | 16.61 | 7.77 |

Supplemental Table S5: Running times of the MATLAB PCA implementation (Vijayan 2020) used in Specter and FIt-tSNE (Linderman et al. 2019) on simulated data. Data sets of different size were simulated using Splatter (same data sets as in Table S4).

| Method | 1k | 10k | 100k | 200k | 500k | 1m |
|---|---|---|---|---|---|---|
| PCA | 0.04s | 0.12s | 0.52s | 1.59s | 3.43s | 5.98s |
| FIt-SNE | 41s | 1m17s | 1m4s | 1m53s | 4m3s | 8m11s |

Supplemental Table S6: Comparison of running times on three largest real data sets. Running times of Specter, Seurat, dropClust, the geometric sketching (Gsketch) based Louvain clustering, TSCAN, and RtsenKmeans are reported in minutes (rounded) on the 3 largest real data sets used in this study. *Running times exclude preprocessing for all methods except TSCAN and dropClust, whose implementation did not allow to isolate the core algorithm. Specter used 50 ensemble members and was run with 20 threads. The last column (Specter+Pre) shows the total running time of Specter and all its preprocessing steps, including log-transformation, selection of highly variable genes (2000), and PCA.

| Data set | #Cells | Specter | Seurat | dropClust* | Gsketch | TSCAN* | RtsneKmeans | Specter+Pre |
|---|---|---|---|---|---|---|---|---|
| CNS | 464,713 | 1 | 11 | 2 | 7 | 3 | 89 | 3 |
| saunders | 665,385 | 2 | 18 | 3 | 19 | 8 | 193 | 4 |
| trapnell | 2,026,641 | 15 | 79 | 12 | 400 | 100 | 1225 | 23 |

### 2 Supplemental Figures

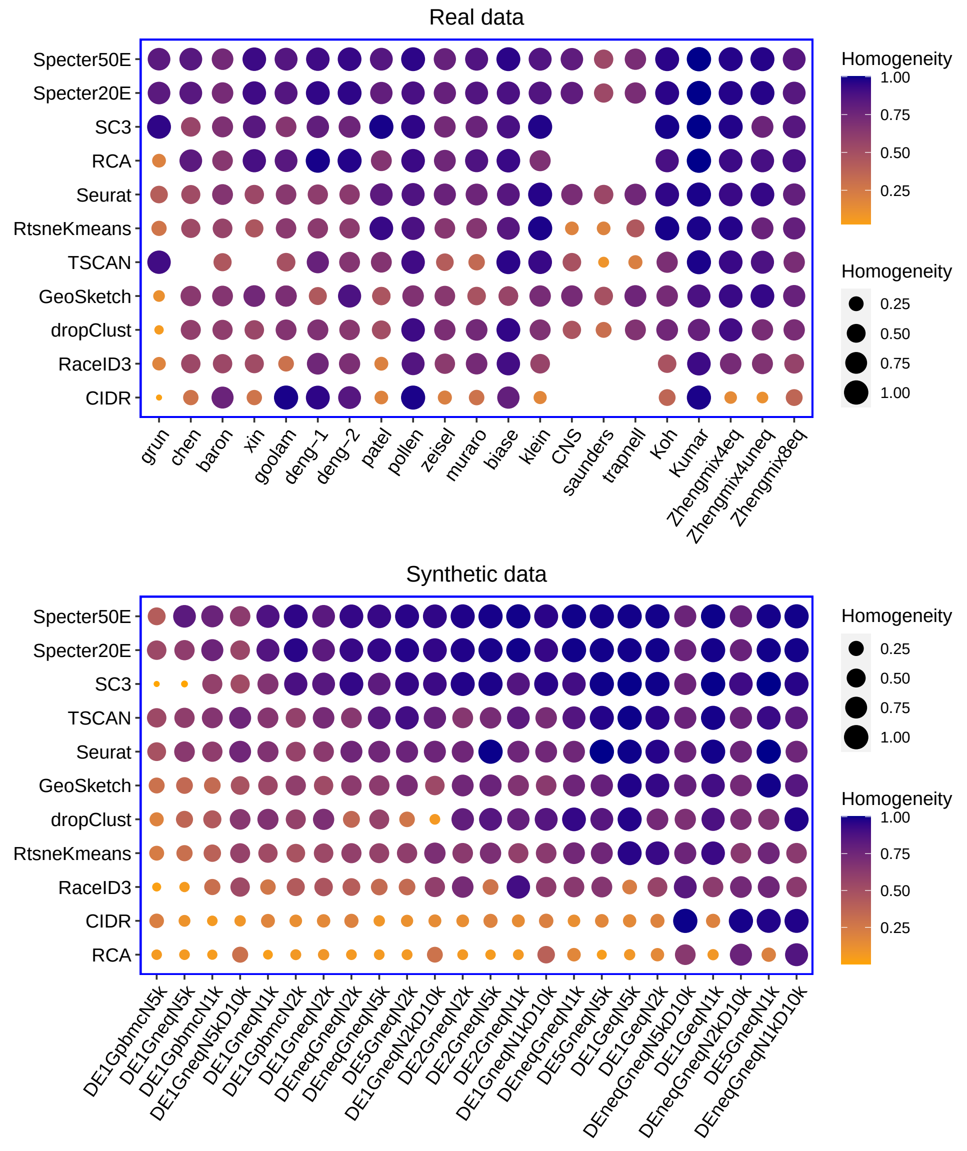


Supplemental Figure S1: Clustering performance measured by homogeneity score of Specter and competing methods on real and synthetic data. Methods are ordered by mean homogeneity score across data sets decreasing from top to bottom. In the calculation of mean scores we excluded for each method the data sets where the method did not run successfully. Synthetic data sets are ordered from left to right by increasing mean homogeneity score over all methods. SC3 , RCA, RaceID3, and CIDR failed to run on the three largest data sets *CNS*, *saunders*, and *trapnell* due to insufficient memory. TSCAN failed to run on data sets *chen* and *skin* for unknown reasons. Geometric sketching refers to the Louvain clustering of 10% of the cells sampled using geometric sketching.

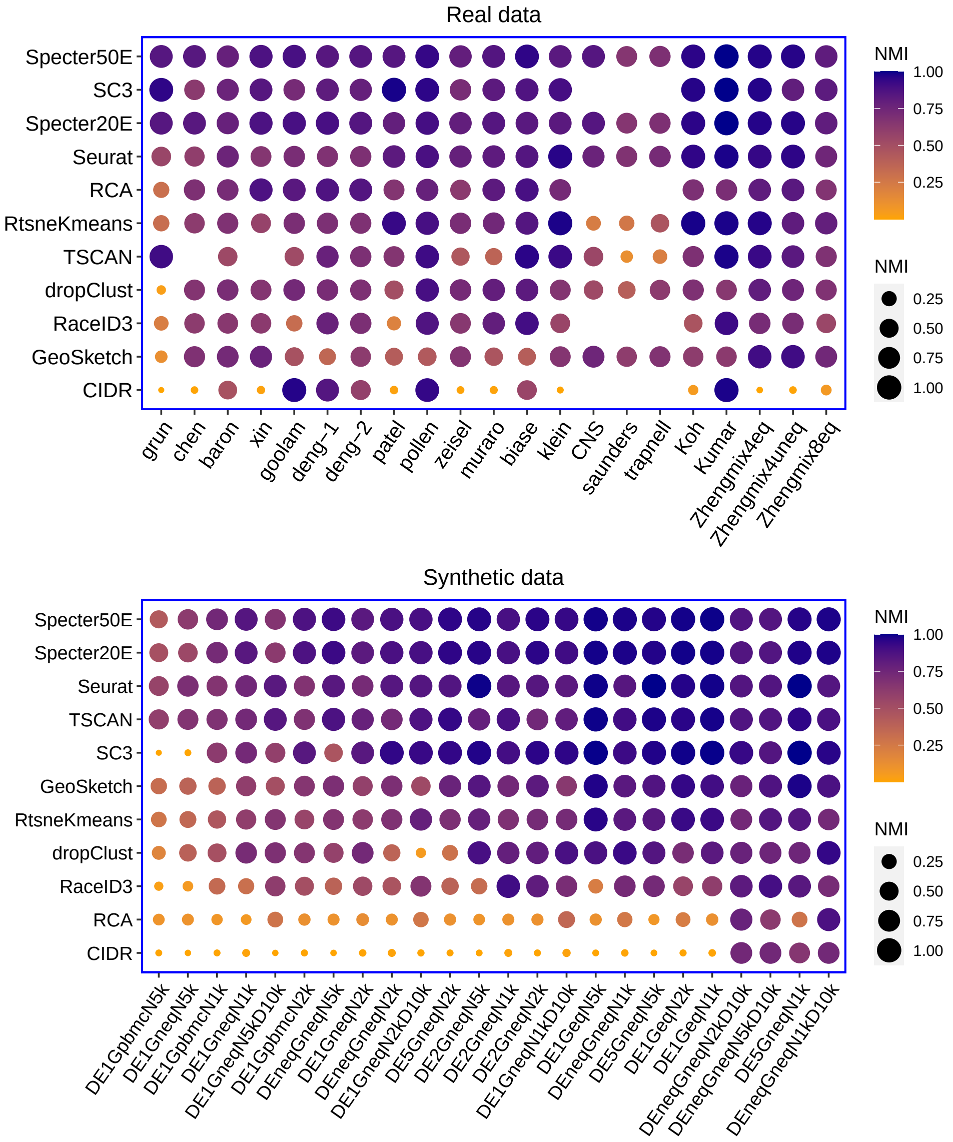


Supplemental Figure S2: Clustering performance measured by NMI of Specter and competing methods on real and synthetic data. Methods are ordered by mean NMI across data sets decreasing from top to bottom. In the calculation of mean scores we excluded for each method the data sets where the method did not run successfully. Restricted to the same set of data sets as SC3, Specter20E was with a mean ARI of 0.87 marginally better than SC3 (mean ARI 0.85). Synthetic data sets are ordered from left to right by increasing mean NMI over all methods. SC3 , RCA, RaceID3, and CIDR failed to run on the three largest data sets *CNS*, *saunders*, and *trapnell* due to insufficient memory. TSCAN failed to run on data sets *chen* and *skin* for unknown reasons. Geometric sketching refers to the Louvain clustering of 10% of the cells sampled using geometric sketching.

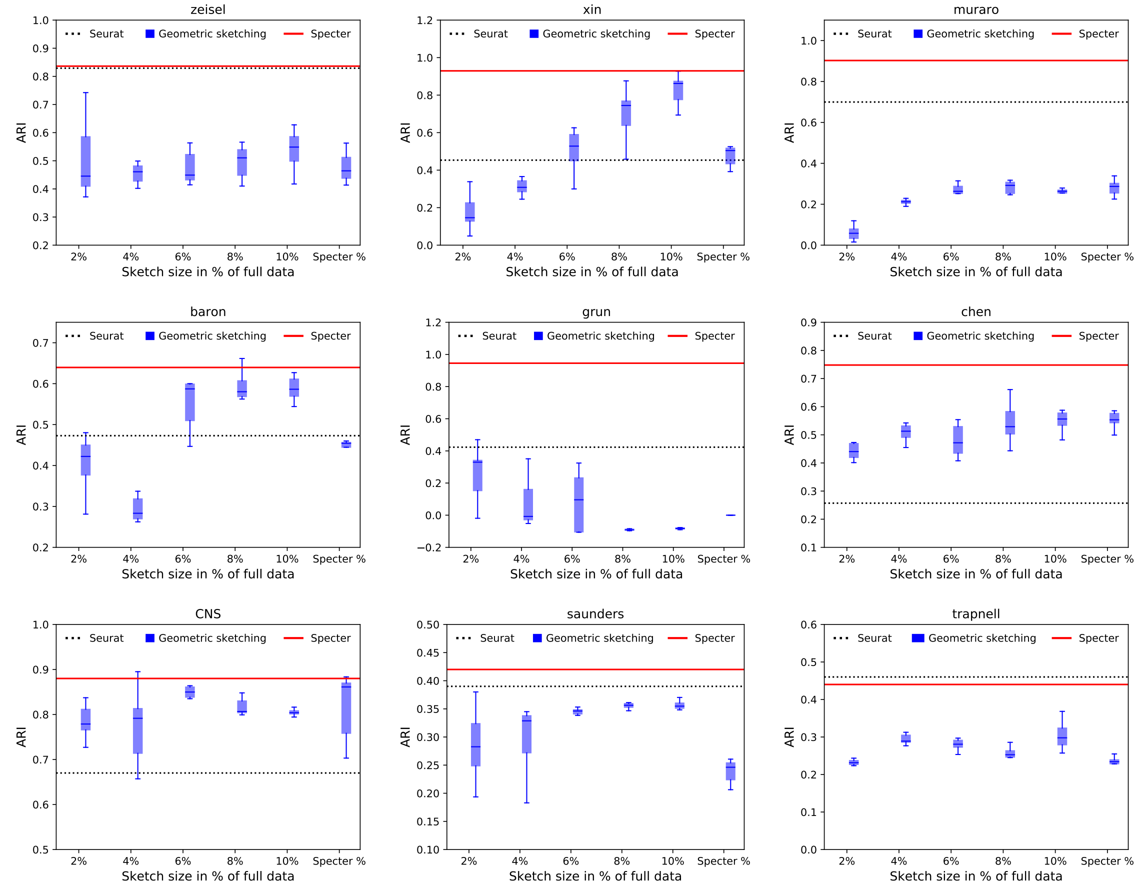


Supplemental Figure S3: Accuracy (in ARI) of geometric sketching based Louvain clustering for varying sketch sizes. For each sketch size, the results of 10 random trials are shown. “Specter %” uses the same number of cells in the geometric sketch as Specter uses landmarks or cells in the selective sampling step (see the “Methods” section), whichever one is larger.

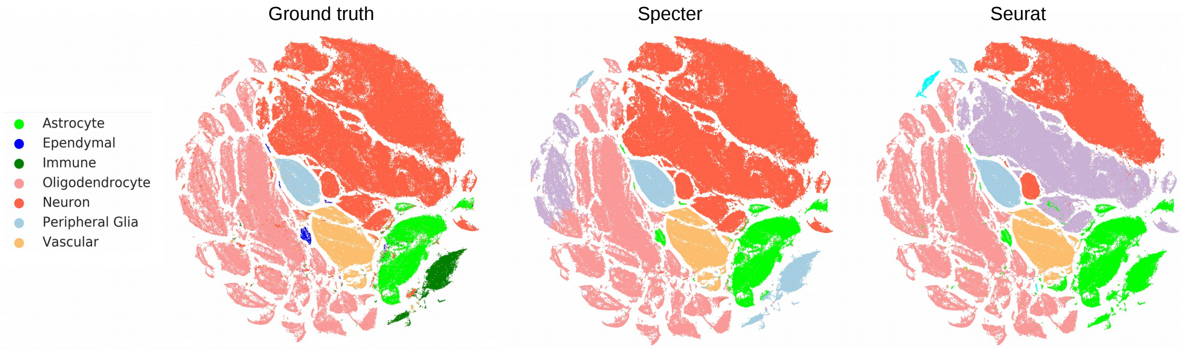


Supplemental Figure S4: t-SNE visualization of single cells of the mouse nervous system (data set *CNS*). Cells in the ground truth representation (left) are colored by cell type specified by the legend. The visualization of Specter (middle) and Seurat (right) clusterings use the same 2D embedding as the ground truth, but cells are colored according to clusters inferred by the two methods; colors do not directly reflect cell types specified by the legend. As expected by the higher ARI (0.89 vs 0.67) (and higher homogeneity scores of 0.81 vs 0.71 and NMI of 0.84 vs 0.78), Specter makes fewer mistakes. In contrast to Specter, Seurat wrongly splits neurons into 2 populations, is not able to distinguish astrocytes from immune cells, and is similarly not able to distinguish a subpopulation of vascular cells from astrocytes.

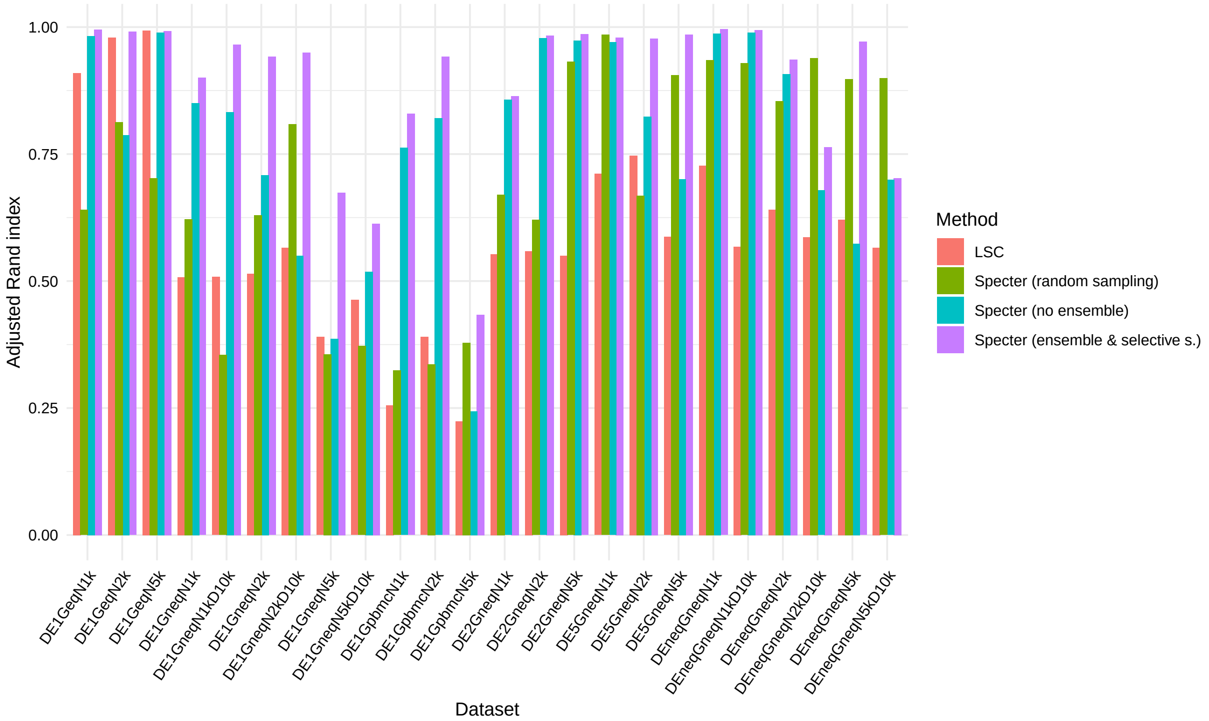


Supplemental Figure S5: Improvements in Specter over LSC. The clustering accuracy of Specter using 50 ensemble members (ensemble & selective s.) is compared to the accuracy of the original implementation of the landmark-based spectral clustering algorithm (LSC) and two variants of Specter in which we either disable consensus clustering in Specter (no ensemble) or in which we replace the novel selective sampling in Specter (Algorithm 2) by random sampling. When no clustering ensemble is used (no ensemble), we set parameters to the median values of intervals probed by the ensemble scheme ($\gamma = 0.15, p = 9k\log(k)$).

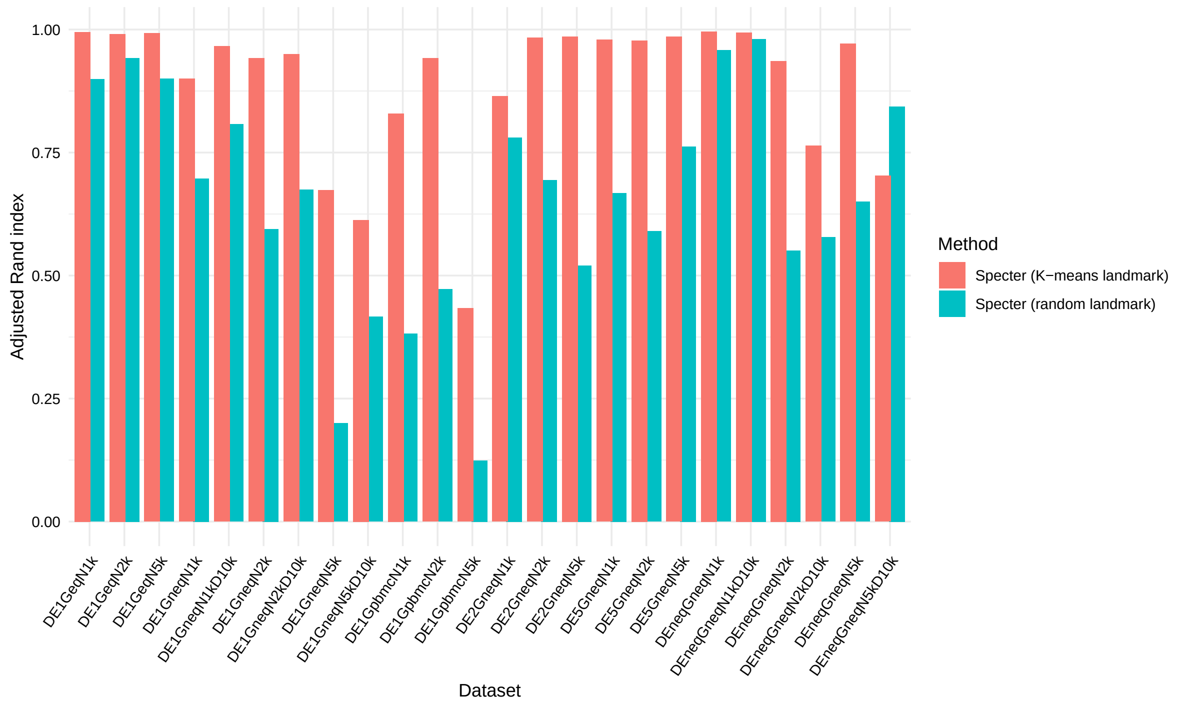


Supplemental Figure S6: Comarison of landmark selection strategies. The clustering accuracy of Specter using our hybrid $k$-means based landmark selection strategy ($K$-means landmark) is compared to a variant of Specter in which we select landmarks uniformly at random.

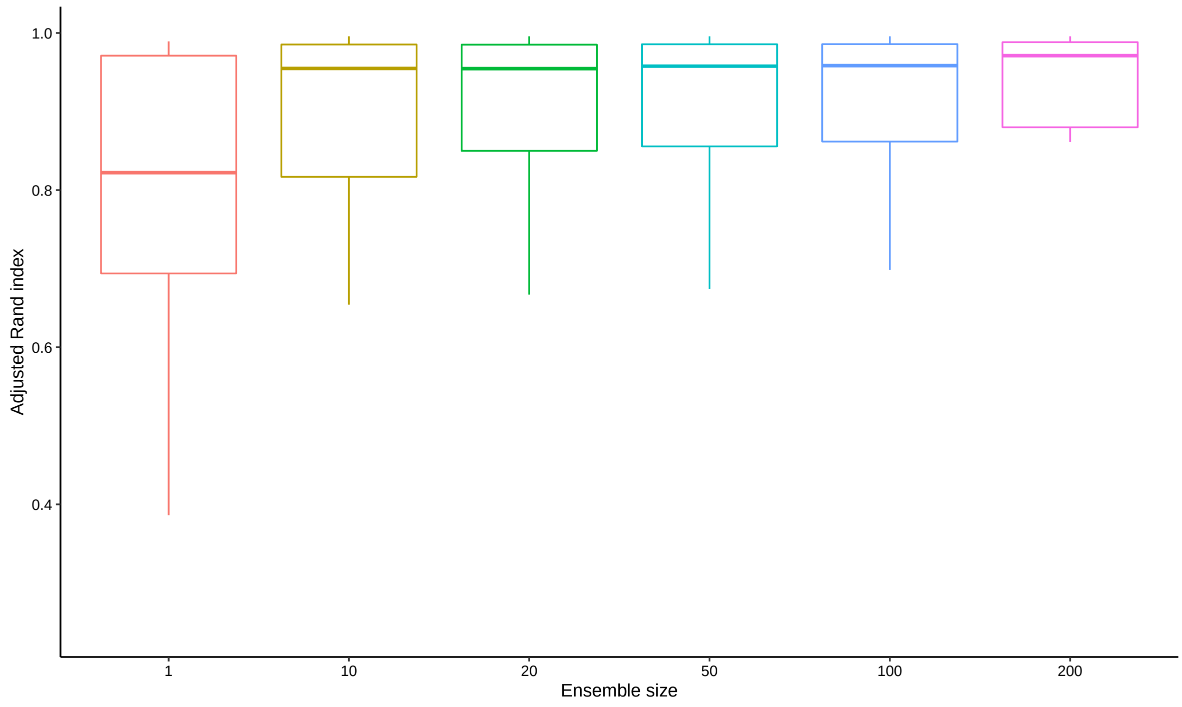


Supplemental Figure S7: Accuracy of Specter vs. number of ensemble members. For each number of ensemble members, the box plot shows minimum, maximum, median, and first and third quartiles of ARI scores achieved by Specter on the 24 simulated data sets described in Table S2.

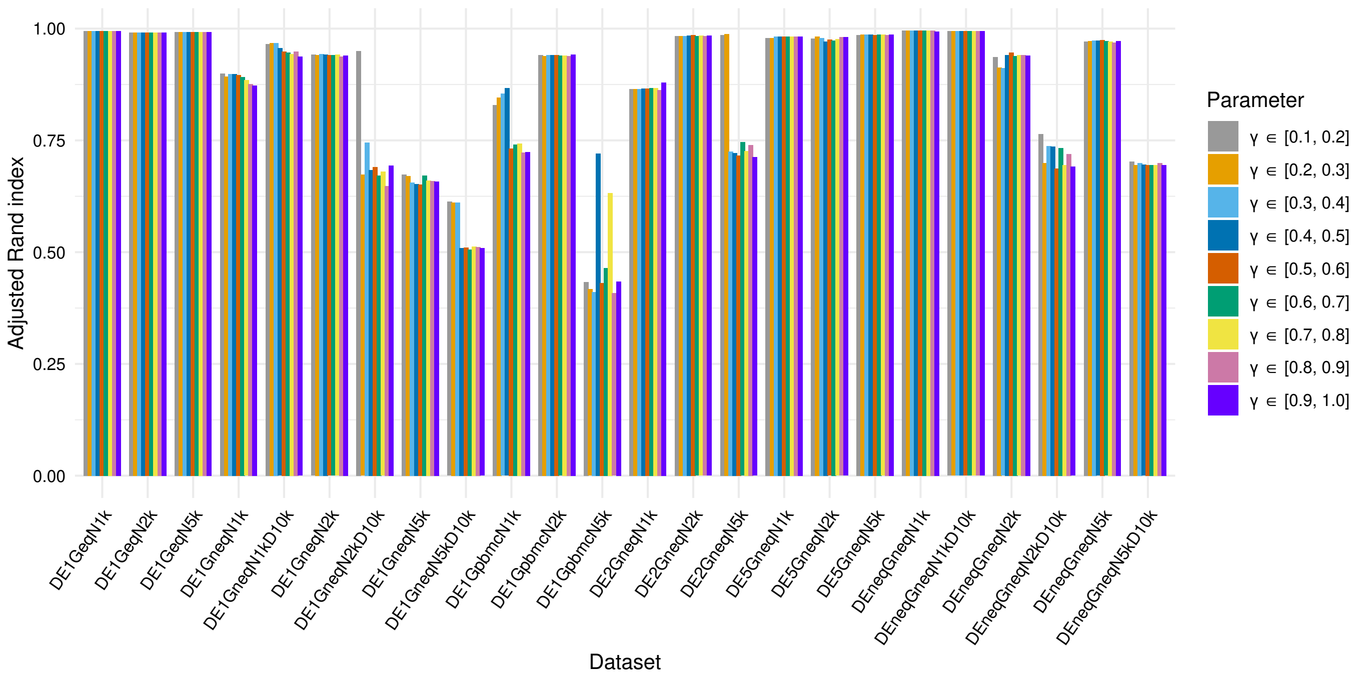


Supplemental Figure S8: Robustness of Specter to choice of parameter $\gamma$. Across 24 synthetic data sets, Specter computed 50 ensemble members using different ranges for parameter $\gamma$. By default, Specter selects a $\gamma \in [0.1, 0.2]$ for each ensemble member.

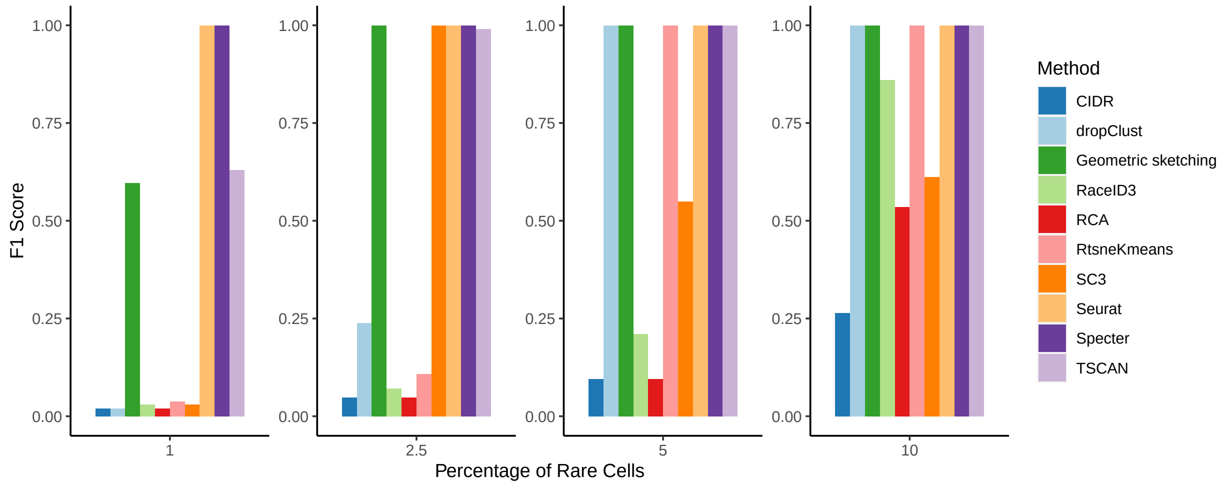


Supplemental Figure S9: Sensitivity to rare cell types with equal starting abundances. 4000 cells from two equal size groups (2000 cells each) were simulated using Splatter. We randomly downsampled one group to comprise 1%, 2.5%, 5%, and 10% of the total number of cells. We repeated this experiment five times for each group and show the average $F_1$ score over the 10 runs. For geometric sketching, the average $F_1$ score was taken over 10 random trials with a sketch size of 10% of the full data.

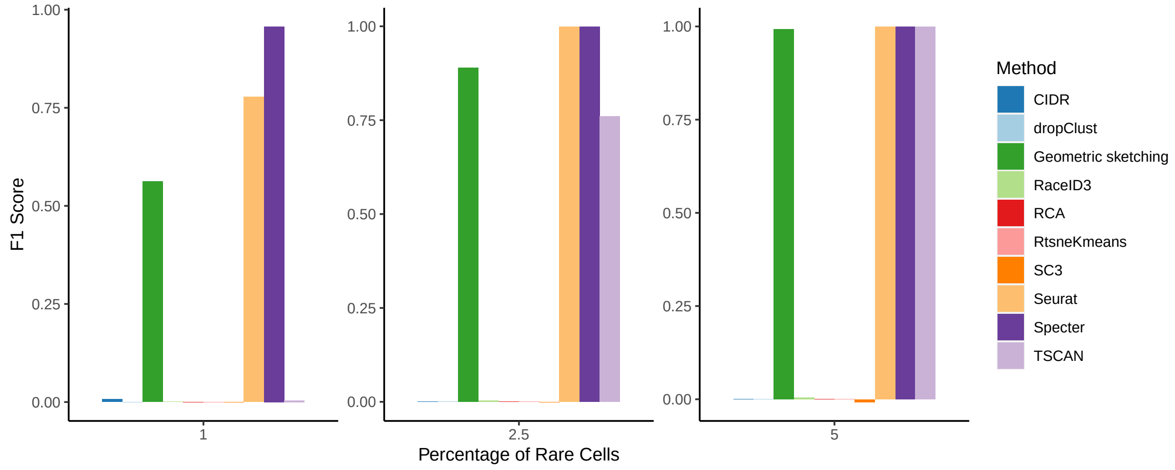


Supplemental Figure S10: Sensitivity to rare cell types in initially smaller group. Cells were randomly sampled from the smaller of two simulated groups (1,000 and 9,000 cells) to comprise 1%, 2.5%, and 5% of the total number of cells. We show the average $F_1$ score over 10 runs of this experiment. For geometric sketching, the average $F_1$ score was taken over 10 random trials with a sketch size of 10% of the full data.

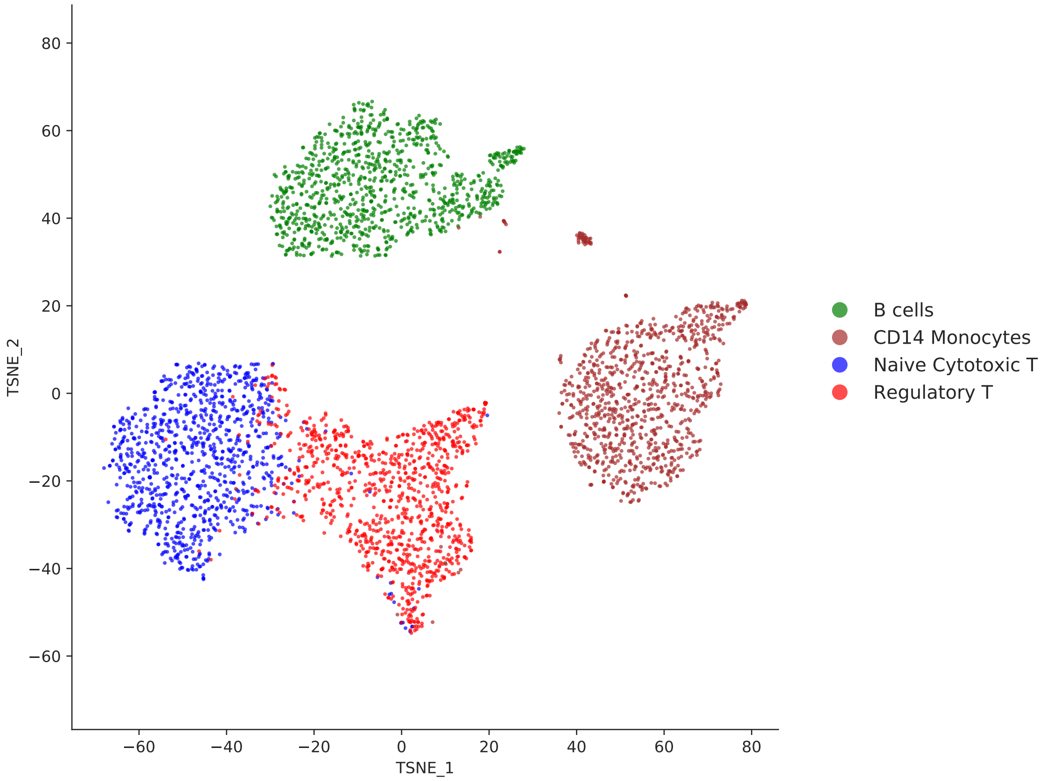


Supplemental Figure S11: t-SNE visualization of the Zhengmix4eq dataset (see Table S1). Naive cytotoxic T cells and regulatory T cells partly overlap.

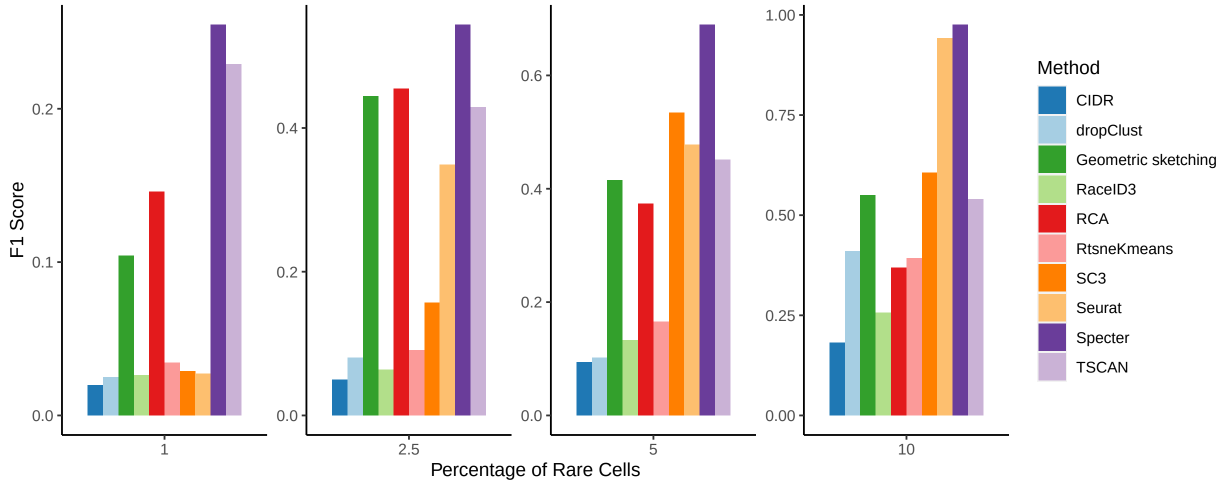


Supplemental Figure S12: Sensitivity to rare population of overlapping cell types. Naive cytotoxic T cells and regulatory T cells taken from the Zhengmix4eq data set overlap in the t-SNE projection shown in Figure S11. We randomly downsampled naive cytotoxic and regulatory T cells to comprise 1%, 2.5%, 5%, and 10% of the total number of cells and repeated this experiment five times for each group. Average $F_1$ scores are shown over the 10 runs, with adjusted $F_1$ score ranges for each subsample size. For geometric sketching, the average $F_1$ score was taken over 10 random trials with a sketch size of 10% of the full data.

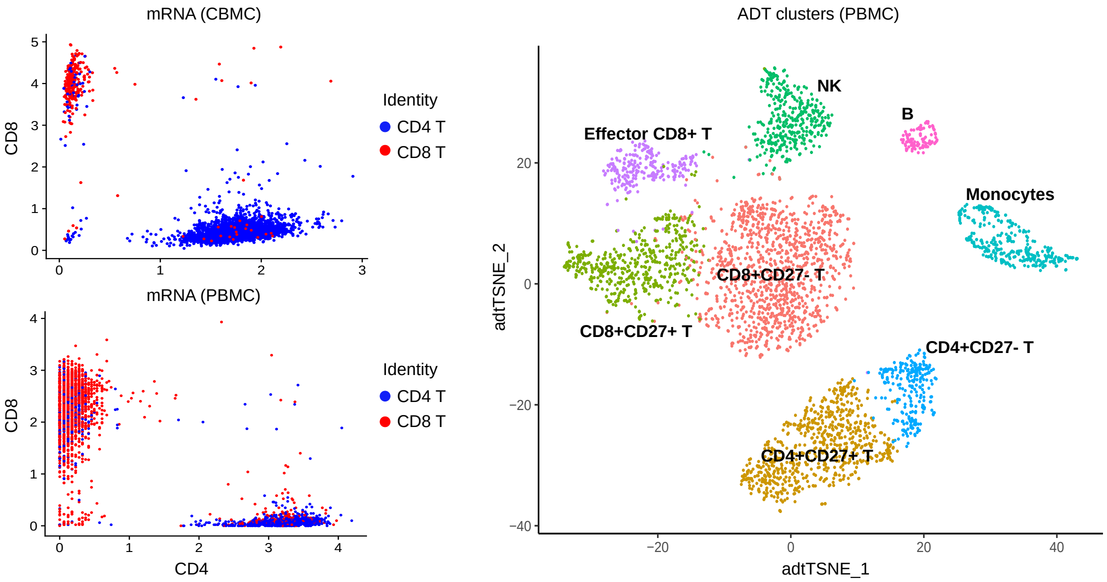


Supplemental Figure S13: Seurat unimodal clustering. *left:* CBMCs (top) and PBMCs (bottom) with coordinates of protein expression (ADT) along CD4 and CD8 axis. Colors denote clusters computed by Seurat based on mRNA expression which contain a mix of CD4 T cells and CD8 T cells. *right:* t-SNE visualization of clusters identified by Seurat from protein expression (ADT) of PBM cells. CD14+ and FCGR3A+ monocytes cannot be discriminated (compare Figure 6), megakaryocytes are not detected.

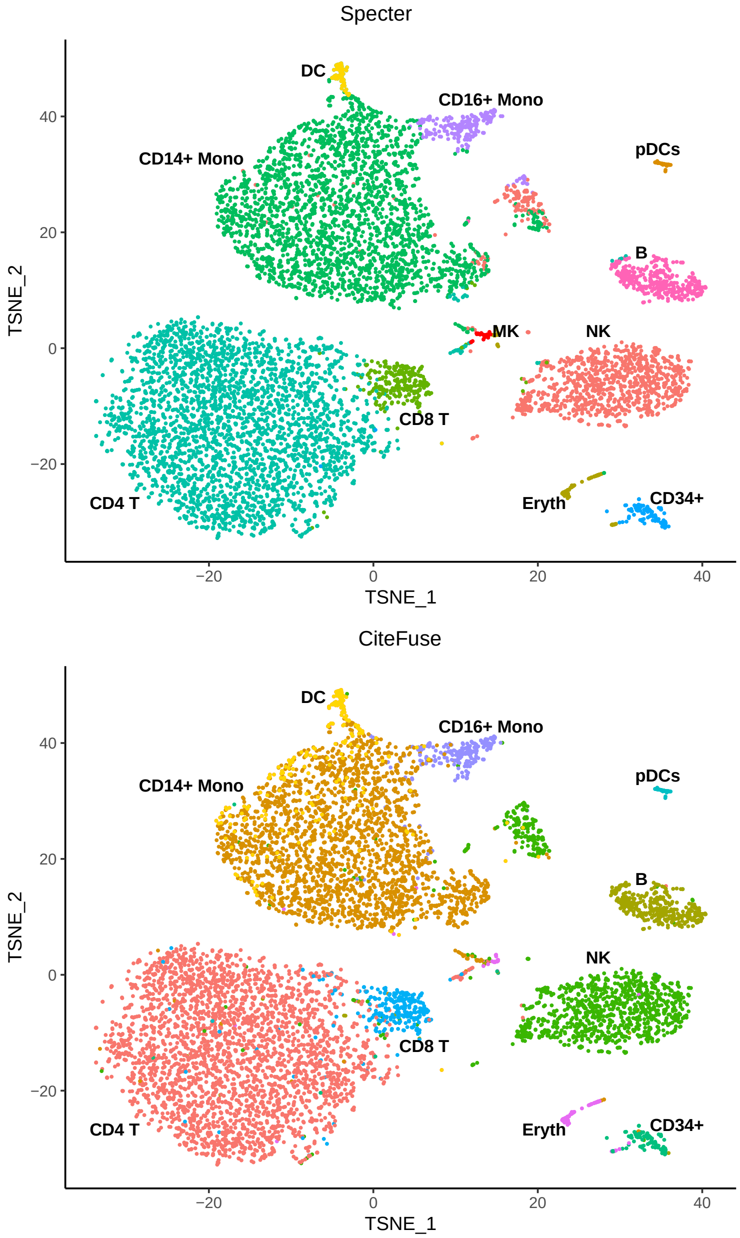


Supplemental Figure S14: Comparison of multimodal clusterings of CBM cells as computed by Specter (top) and CiteFuse (bottom). Despite an overall high agreement between the two clusterings (ARI 0.94), only Specter detects a rare population of megakaryocytes (red).

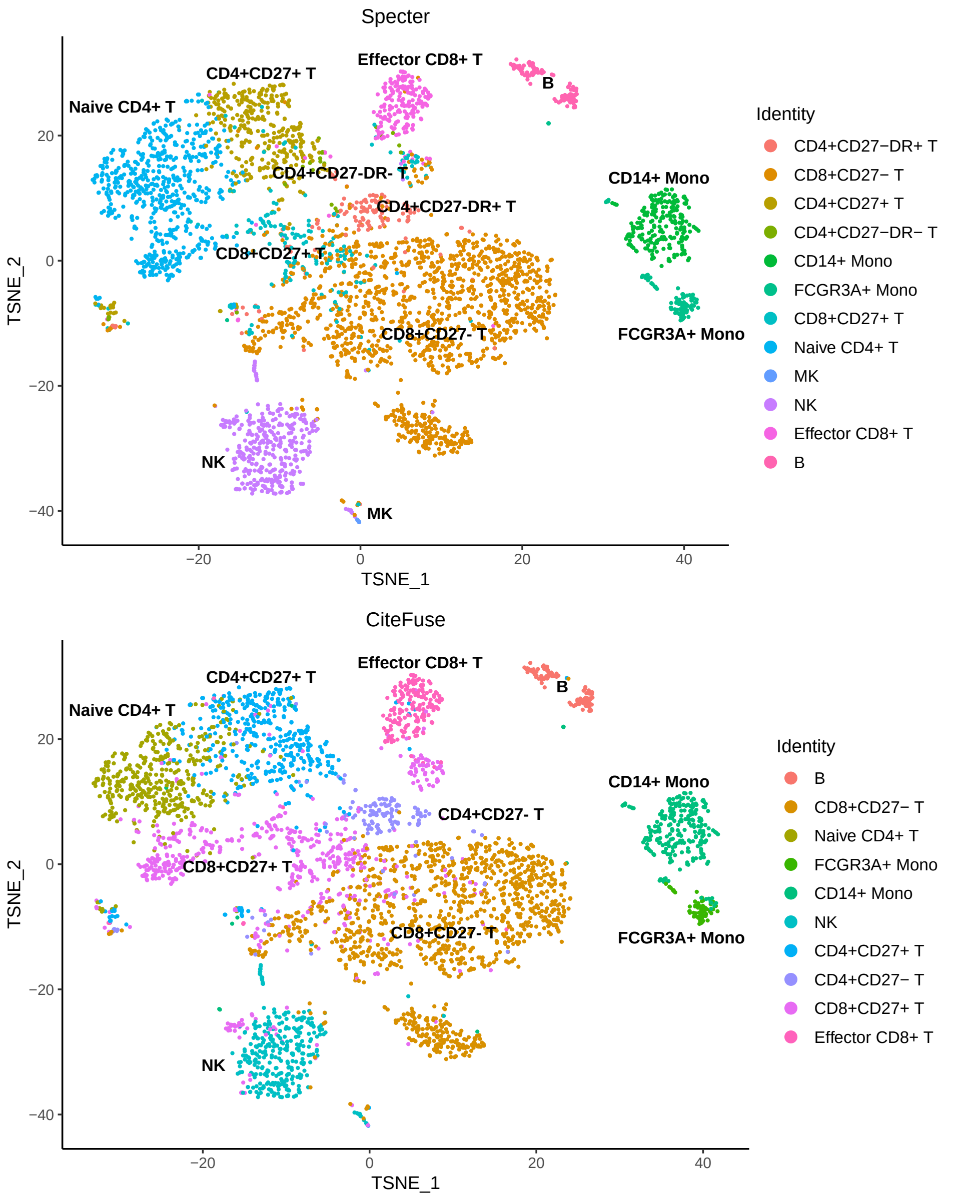


Supplemental Figure S15: Comparison of multimodal clusterings of PBM cells as computed by Specter (top) and CiteFuse (bottom). Despite an overall high agreement between the two clusterings (ARI 0.86), only Specter detects a rare population of megakaryocytes and can discriminate between CD27-DR+ and CD27-DR- subpopulations of CD4+ memory T cells.

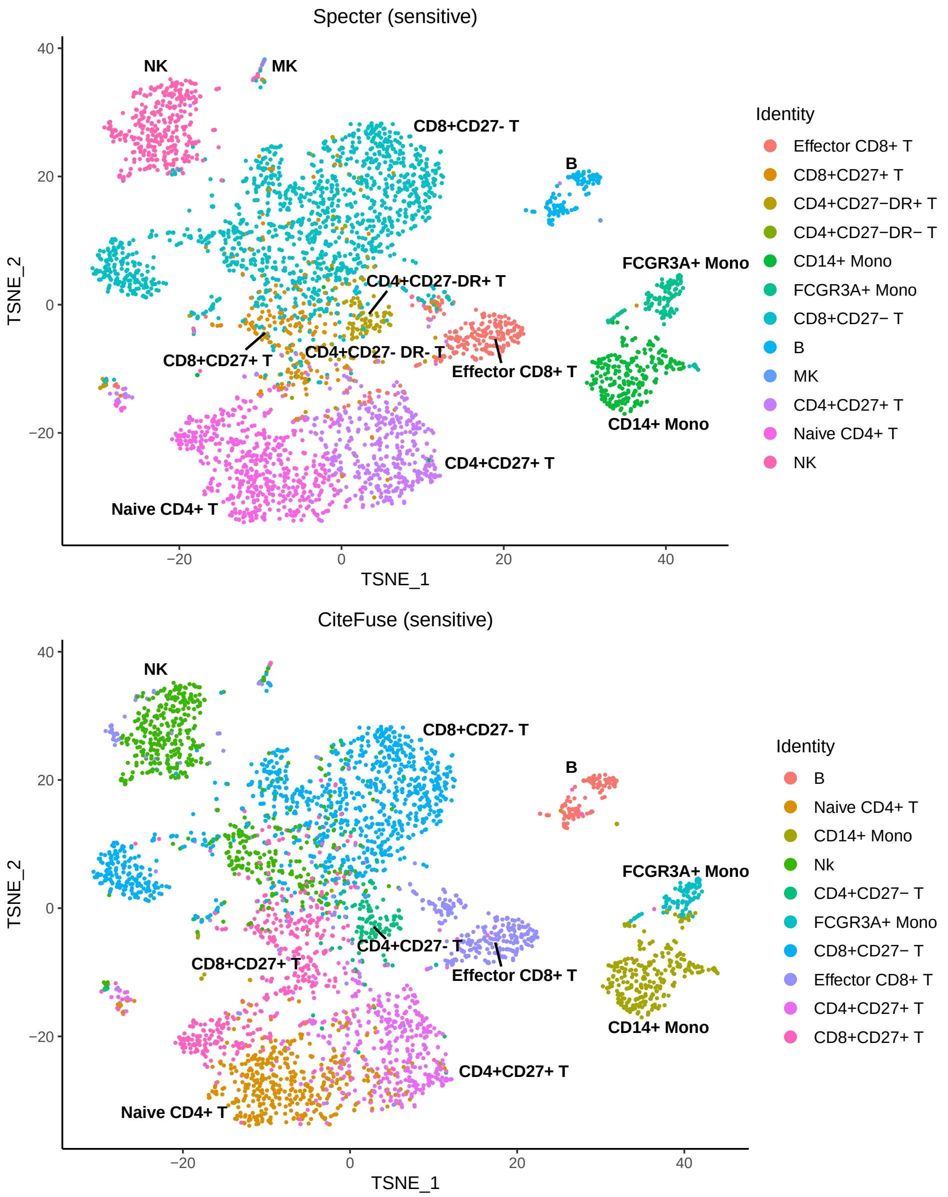


Supplemental Figure S16: Comparison of multimodal clusterings of PBM cells. Here, Specter (top) and CiteFuse (bottom) use slightly more conservative parameters in the doublet removal (eps = 190, minPts = 10). Again, only Specter is able to discriminate between CD27-DR+ and CD27-DR- subpopulations of CD4+ memory T cells and detects a rare population of megakaryocytes.

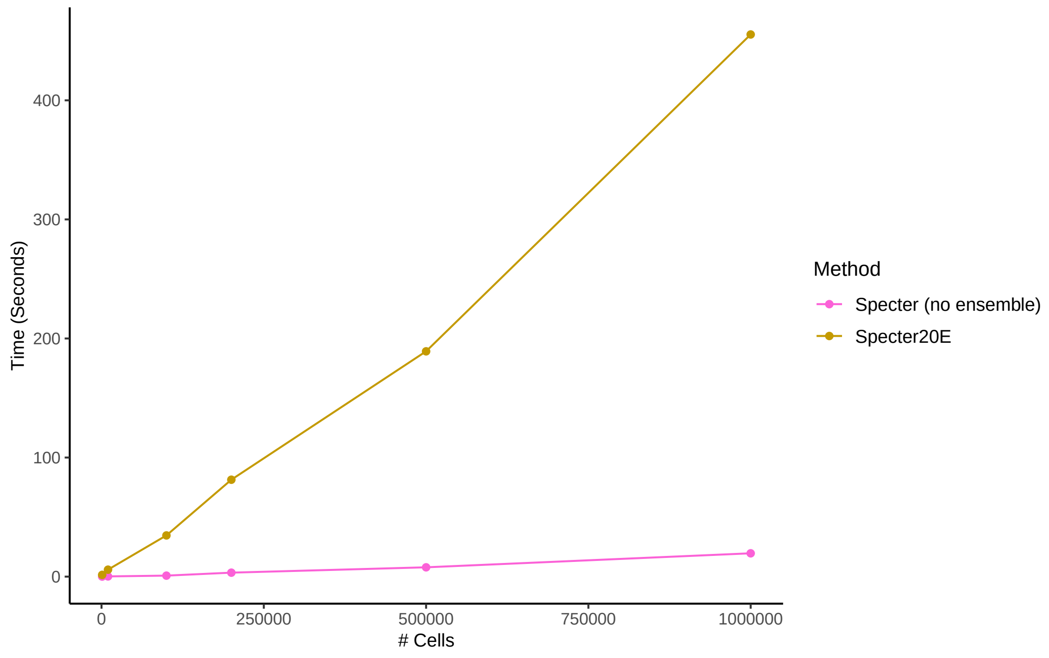


Supplemental Figure S17: Linear-time complexity of Specter. CPU times in seconds (single threaded) are shown for the core algorithm of Specter (no ensemble) and Specter using a clustering ensemble of size 20. Different size data set were simulated using Splatter containing 1k, 10k, 100k, 200k, 500k, and 1 million cells.

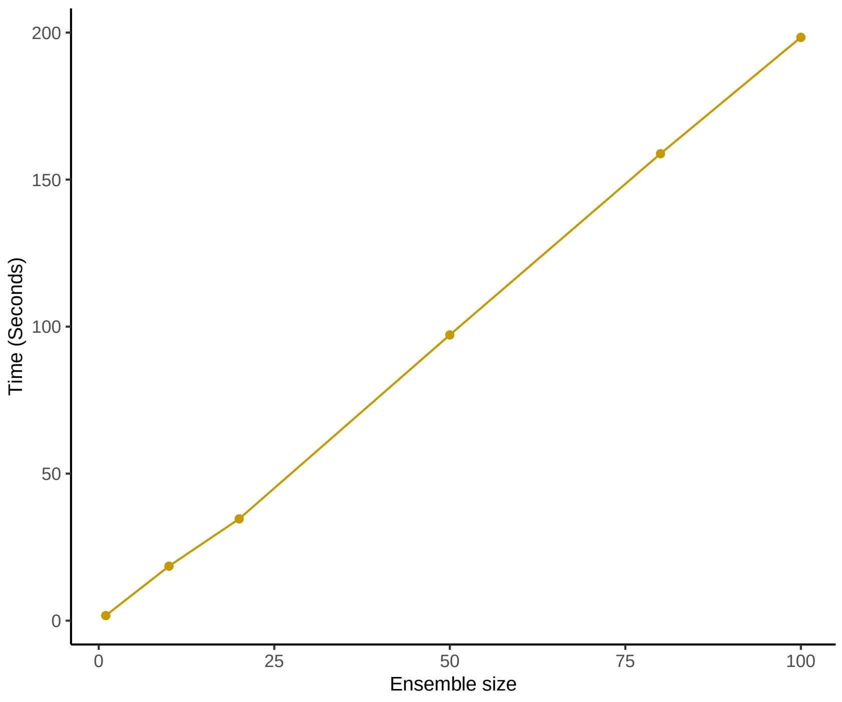


Supplemental Figure S18: Linear increase in running time with number of ensemble members. CPU times in seconds (single threaded) are shown for Specter using an increasing number of ensemble members on a simulated data set containing 100,000 cells.

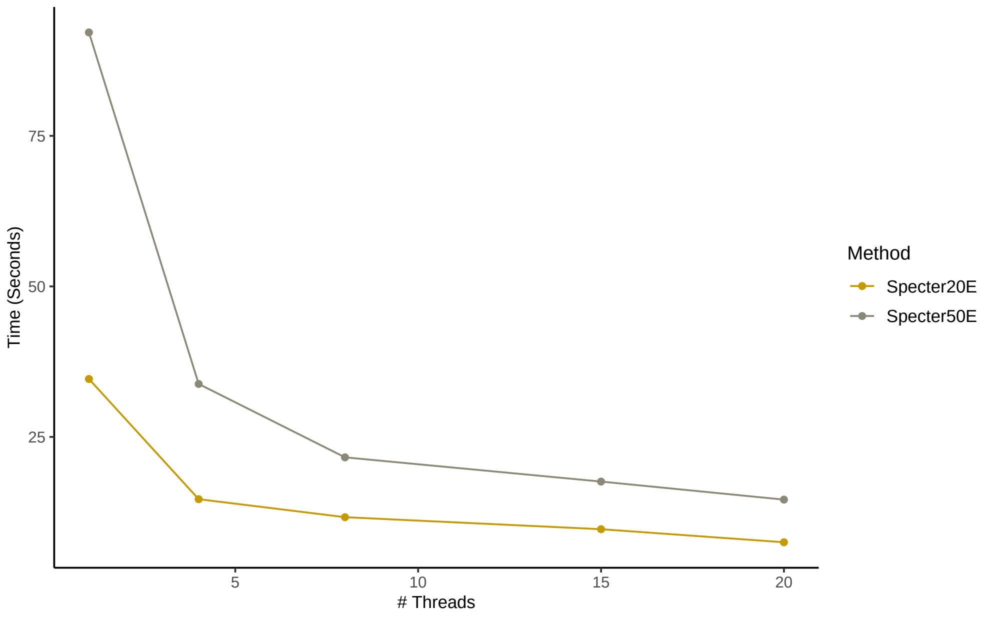


Supplemental Figure S19: Specter speed-up with number of threads. CPU times in seconds are shown for Specter using an increasing number of threads on a simulated data set containing 100,000 cells. 20 or 50 clustering ensemble members were used.

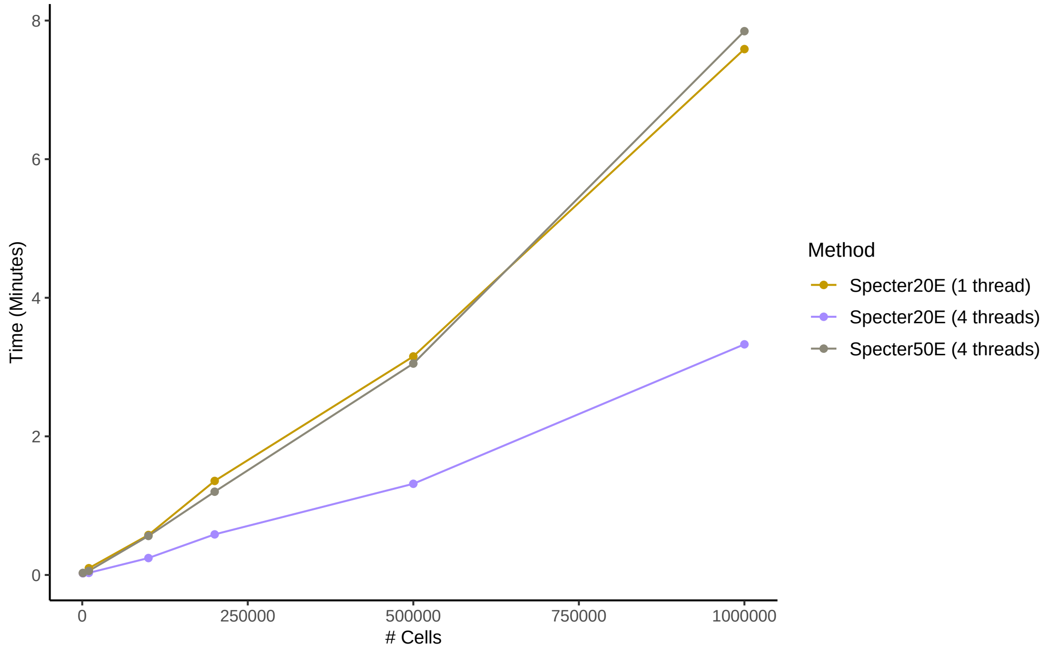


Supplemental Figure S20: Increase in running time for fixed number of threads. CPU times in minutes are shown for Specter using 20 or 50 clustering ensemble members and 1 or 4 threads.

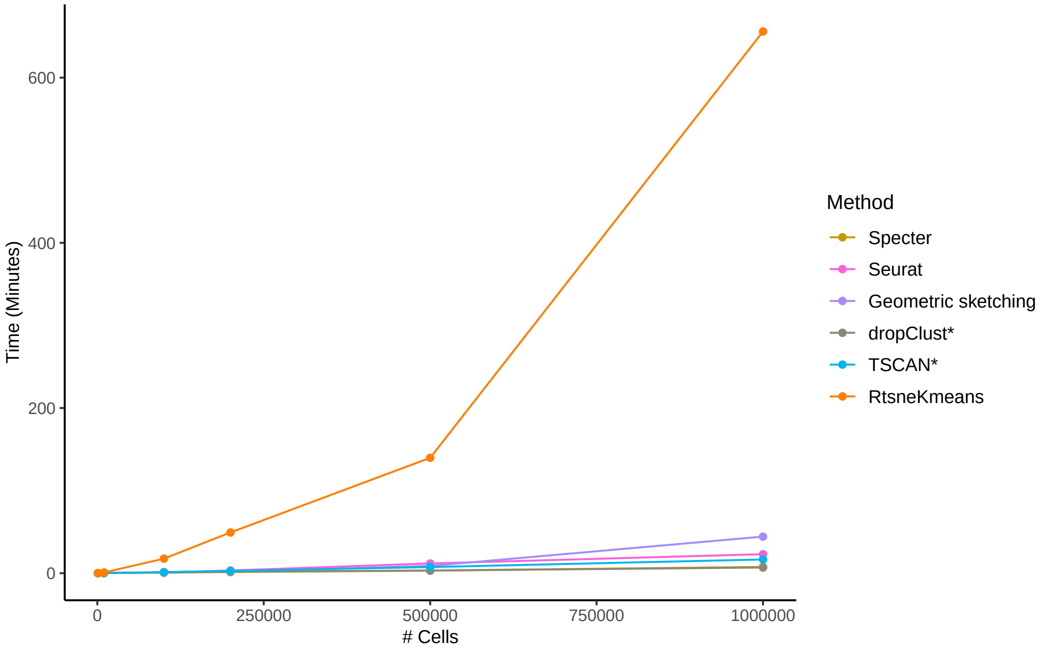


Supplemental Figure S21: Runtime comparison between methods as a function of sample size. CPU times are shown in minutes on different numbers of cells sampled from a simulated data set containing 1 million cells. Seurat was run with a call to the more efficient SCANPY implementation of the Louvain clustering algorithm. *Running times exclude preprocessing for all methods except TSCAN and dropClust, whose implementation did not allow to isolate the core algorithm.
